# Interconnected reservoirs of multidrug-resistant bacteria and plasmids within hospital plumbing infrastructure

**DOI:** 10.64898/2026.07.20.739580

**Authors:** Medini K. Annavajhala, Todd W. Hokunson, Kasiani Terzoglou, Anne-Catrin Uhlemann, Angela Gomez-Simmonds

## Abstract

Hospital sinks and wastewater streams are hotspots for contamination and colonization with multidrug-resistant Gram-negative bacteria (MDR-GNB). While hospital water systems may facilitate MDR-GNB transmission during outbreaks, longitudinal population dynamics of environmental MDR-GNB and their long-term relationship with patient isolates remain unclear. Here we aimed to characterize the diversity and abundance of MDR-GNB within the hospital built environment and evaluate their relationship with clinical isolates. We performed nanopore sequencing on MDR-GNB cultured from sink drains (n=523) and wastewater (n=146) at a New York City hospital between March-August 2023. We assessed spatial and longitudinal genomic strain and plasmid dynamics and quantified relatedness to historical clinical isolates from the same hospital (n=602). Sink drains (54-73%) and wastewater (100%) demonstrated widespread, longitudinal colonization with MDR-GNB, driven primarily by carbapenem-resistant GNB including opportunistic organisms and Enterobacterales encoding *bla*_KPC_. While bacterial colonization differed across environmental niches, we identified both MDR-GNB strains and resistance plasmids that were shared between clinical and environmental sources, as well a distinct interplay between clinical and wastewater versus sink drain samples. This study supports the role of hospital plumbing as an important reservoir for clinically relevant MDR-GNB, but highlights differences across environmental sources, which may impact surveillance approaches.

## Introduction

Infections caused by antibiotic-resistant bacteria cause over 2.8 million infections per year in the US and are associated with disproportionately high morbidity, mortality, and healthcare costs.^1^ Among the most urgent and serious threats to human health identified by the Centers for Disease Control and Prevention are multidrug-resistant Gram-negative bacteria (MDR-GNB) including carbapenem-resistant and extended-spectrum beta-lactamase (ESBL)-producing Enterobacterales (CRE, ESBLE).^1^ MDR-GNB are responsible for a high burden of infections in healthcare settings,^2^ where they have a propensity to spread both directly through patient contact and indirectly through the hospital environment.^3,4^ Importantly, findings from recent genomic studies indicate that indirect transmission of MDR-GNB to patients is difficult to detect and control using traditional infection control methods,^5,6^ signaling a need to better understand environmental reservoirs of MDR-GNB to improve surveillance and mitigation strategies.

Within the hospital environment, hospital water systems have the potential to harbor a wide diversity of GNB, including both enteric bacteria and environmental organisms associated with opportunistic infections in humans.^7^ Hospital sinks and drains are established sites for contamination with MDR-GNB along with other water-containing equipment and fixtures such as toilets, basins, and showers, which are ubiquitous in patient care settings.^8,9^ Longitudinal genomic studies have found strong evidence for sink-to-patient transmission of MDR-GNB,^10,11^ which is thought to be driven by droplet dispersion of resident bacteria from contaminated drains.^12^ Drains and sink traps are difficult to disinfect and may provide the ideal microenvironment for bacterial colonization and biofilm formation, facilitating long-term MDR-GNB persistence.^13^ Sink and drain contamination may also contribute to the dissemination of MDR bacteria into hospital grey water and wastewater streams. However, the relationship between bacterial populations found in distinct components of hospital water distribution and plumbing infrastructure requires further study.

While clonal transmission is a frequent target of surveillance and infection control efforts, plasmid-mediated transmission is increasingly recognized as a widespread contributor to the dissemination of multidrug resistance in hospitals.^14–17^ Hospital water systems have been suggested as important reservoirs for plasmids and other mobile genetic elements (MGEs) harboring antibiotic resistance genes (ARGs), which contribute to ARG transmission and evolution of MDR bacteria via horizontal gene transfer (HGT).^18^ In prior analyses of plasmids harboring the carbapenemase gene *bla*_KPC_, our group identified highly conserved plasmids in clinical cultures collected over many years, suggesting the presence of persistent, hospital-based plasmid reservoirs.^14,19–21^ This is supported by several genomic studies demonstrating putative plasmid sharing between environmental and patient bacterial strains as well as ongoing MDR-GNB colonization and possible plasmid exchange within the hospital environment and wastewater streams.^6,22–24 14–17HGT^ may also represent an important means for plasmid persistence by enabling the transmission of plasmids between human pathobionts and bacteria well-adapted to survive in the environment.

Here we investigated the genomic epidemiology of MDR-GNB isolated from the hospital environment to identify putative interactions between bacteria isolated from clinical cultures, hospital unit sink drains, and hospital wastewater. We hypothesized that the MDR-GNB colonizing hospital environmental sites are highly interconnected by the exchange of specific bacterial lineages and ARG-harboring plasmids and reflect infections in collocated patient populations. By exploring these links, we aimed to better understand how the environment contributes to the nosocomial persistence and spread of MDR-GNB in an urban center with a high burden of antibiotic resistance.^25^ We used a combination of bacterial culture and nanopore whole-genome sequencing to isolate and characterize MDR-GNB from the hospital environment, including analysis of fully resolved plasmids harboring the antibiotic resistance genes *bla*_KPC,_ *bla*_NDM,_ and *bla*_CTX-M_, and quantified chromosome and plasmid sharing between environmental and clinical isolate collections. We found surprisingly little evidence for common MDR-GNB signatures across environmental sample types, reflecting niche-specific bacterial communities within the hospital plumbing infrastructure. Despite this, we identified significant sharing of chromosomes and resistance plasmids between clinical and environmental isolates, supporting hospital water sources as important reservoirs for clinically relevant MDR-GNB and highlighting the unique roles of potential transmission routes in shaping MDR-GNB dynamics within hospital environments and patient populations. Ultimately, these findings can inform and help shape comprehensive environmental surveillance and novel infection control approaches to mitigate the clinical impact of MDR-GNB.

## Results

### Environmental MDR-GNB prevalence and persistence

To assess the overall burden and diversity of MDR-GNB in hospital sink drains, we implemented five timepoints for sink drain sampling between March and September 2023 (TP1-TP5) in two inpatient hospital units (inpatient hospital unit (IHU) A and IHUB) (**Figure 1**). IHUA and IHUB each contain a total of 37 sink drains, including the sink in each unit’s hospital staff bathroom (**Figure S1A**).

**Figure 1.**
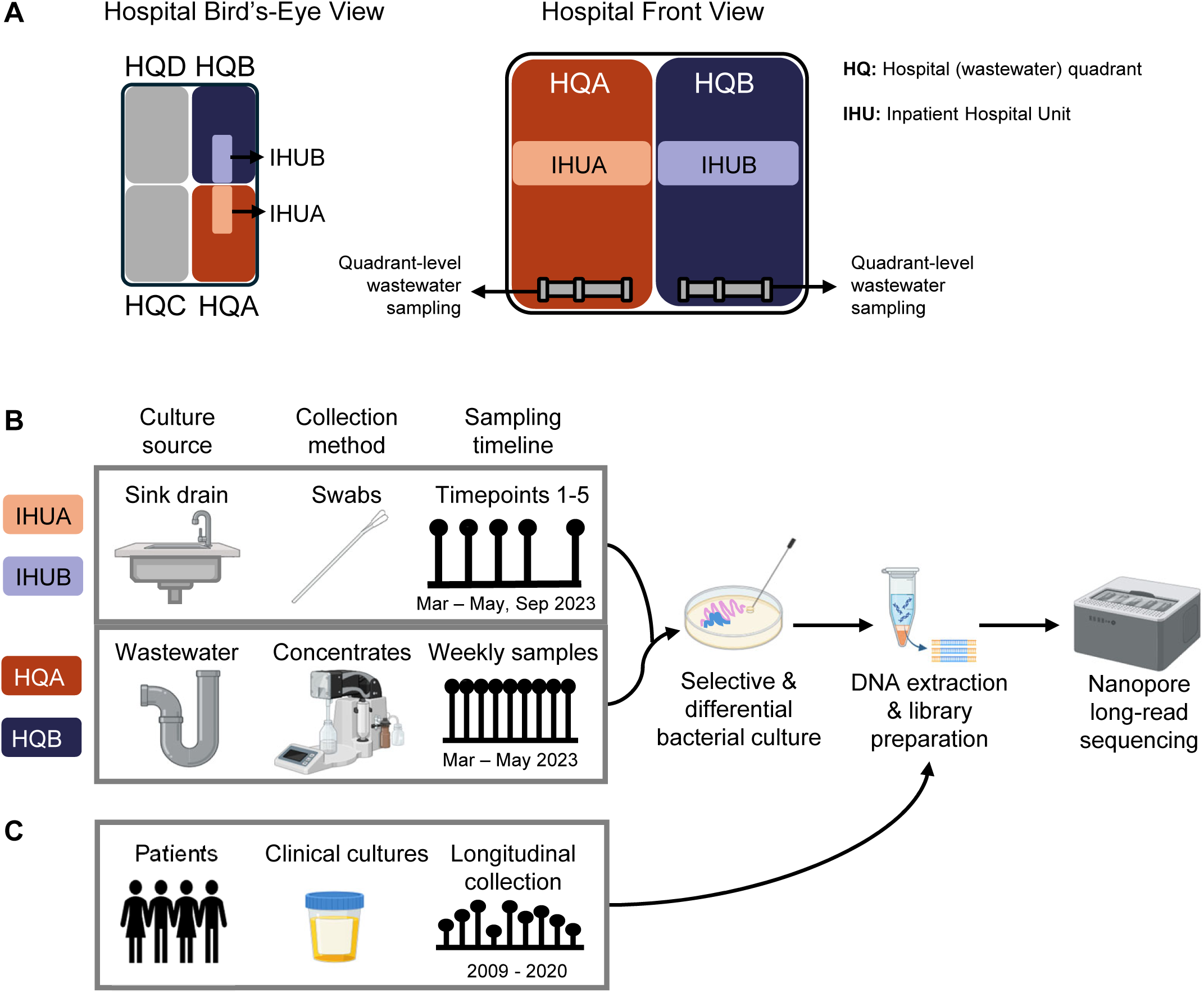
Environmental Surveillance Sampling Strategy. (A) Bird’s eye (left) and front-facing (right) representations of hospital building sampling setup. The hospital plumbing infrastructure is comprised of four vertical hospital quadrants (HQ); all wastewater and grey water streams from a given quadrant flows through one of four house traps prior to entry into the municipal system (HQA, HQB, HQC, HQD). Sampling of HQA and HQB house traps was implemented as part of broader efforts related to viral and bacterial pathogen surveillance in hospital wastewater. Inpatient hospital units A and B (IHUA, IHUB) are adjacent units on the same floor of the hospital. Each unit corresponds to a different vertical plumbing quadrant; i.e., wastewater from IHUA flow through the HQA house trap along with wastewater from other floors in quadrant A, and same for IHUB and the HQB house trap. **(B)** Overall sampling strategy and timeline for this project. All accessible sink drains in patient rooms and staff unit bathrooms in IHUA and IHUB were sampled using culture swabs over five timepoints, roughly monthly from March to May 2023 and again in September 2023. HQA and HQB house trap wastewater was sampled weekly, and microbes were concentrated from raw wastewater. The full wastewater collection ranged from January 2022 – July 2023; samples from date ranges overlapping with sink drain swabbing timepoints in March to May 2023 were selected for analysis in this study. All sink drain and wastewater samples underwent selective and differential bacterial culture to isolate diverse multidrug-resistant Gram-negative bacteria (MDR-GNB). DNA was then extracted from each isolate and underwent long-read nanopore sequencing. **(C)** We took advantage of a longitudinal collection of clinical cultures from patients within the hospital system, including but not limited to IHUA and IHUB, which encompassed MDR bacterial isolates collected from 2009 through 2020 as part of routine clinical care. Clinical isolates underwent long-read sequencing for comparison to environmental isolate genomes in this study. Figure icons were sourced from BioRender with institutional licensing and the NIH NIAID BIOart resource. ***Abbreviations***: HQ – hospital (wastewater) quadrant; IHU – inpatient hospital unit

We sampled 62-92% of available sinks in IHUA and 68-86% of available sinks in IHUB at each timepoint. Overall, MDR-GNB were highly prevalent, with 50-74% of sampled sinks in IHUA and 71-90% of sampled sinks in IHUB identified as MDR-GNB-positive (MDR+) based on selective culture (**Figure S1**). Carbapenem-resistant (CR)-GNB were the primary drivers of sink drain MDR-GNB colonization, identified in 92% of both IHUA and IHUB sinks, as compared to 3^rd^ generation cephalosporin-resistant (CephR)-GNB (41% of IHUA and 51% of IHUB sink drains). A higher proportion of sink drains were also persistently colonized over two or more consecutive timepoints with CR-GNB (54% of IHUA and 73% of IHUB sink drains) than Ceph-RE (14% of IHUA and 22% of IHUB sink drains; **Figure S2**). However, across individual sinks, CR- and CephR-GNB detection were significantly associated (χ^2^ (df=1, N=297)=5.8, p=0.02; **Figure S3**). We separately collected wastewater samples weekly as part of broader pathogen surveillance efforts from hospital quadrants HQA and HQB,^26^ which receive wastewater from inpatient hospital units IHUA and IHUB, respectively (**Figure 1**), during concomitant timepoints TP1-TP4. Wastewater from both HQA and HQB was universally positive for MDR-GNB, including both CephR-GNB and CR-GNB, highlighting their ubiquity in hospital wastewater and grey water discharge systems (**Figure S1B**).

In total, 669 MDR-GNB isolates were recovered from 297 sink drain swabs and 36 wastewater samples across the project timeline. After excluding isolates collected from the same environmental sample with identical sequence type (ST) and ARG profiles to mitigate bias due to our selective culture and colony isolation protocol, we analyzed 443 total MDR-GNB genomes (**Table 1**).

**Table 1.** Unique multidrug-resistant gram-negative bacterial isolates collected from each environmental site across sampling timepoints.

| COLLECTION SITE | ISOLATE PHENOTYPE<sup>1</sup> | TP1 ISOLATES | TP2 ISOLATES | TP3 ISOLATES | TP4 ISOLATES | TP5 ISOLATES<sup>2</sup> | TOTAL ISOLATES |
|---|---|---|---|---|---|---|---|
| **IHUA** | CephR | 2 | 7 | 5 | 8 | 4 | 26 |
|  | CR | 26 | 28 | 26 | 21 | 31 | 132 |
|  | Total MDR | 28 | 35 | 31 | 29 | 35 | 158 |
| **IHUB** | CephR | 6 | 10 | 5 | 17 | 4 | 42 |
|  | CR | 45 | 27 | 28 | 26 | 30 | 156 |
|  | Total MDR | 51 | 37 | 33 | 43 | 34 | 198 |
| **HQA** | CephR | 9 | 4 | 3 | 4 | - | 20 |
|  | CR | 10 | 10 | 3 | 3 | - | 26 |
|  | Total MDR | 19 | 14 | 6 | 7 | - | 46 |
| **HQB** | CephR | 8 | 6 | 2 | 5 | - | 21 |
|  | CR | 7 | 5 | 4 | 4 | - | 20 |
|  | Total MDR | 15 | 11 | 6 | 9 | - | 41 |

<sup>1</sup> As determined by selective culture. CephR = third-generation cephalosporin-resistant; CR = carbapenem-resistant. CephR phenotype defined as growth on CHROMagar ESBL media and absence of growth on CHROMagar KPC media; CR phenotype defined as growth on CHROMagar KPC media.
<sup>2</sup> Wastewater isolates only collected during timepoints TP1-TP4 as described in methods.

### Environmental niche-specificity of MDR-GNB

MDR-GNB isolated from sink drains and wastewater were largely distinct across collection sites (IHUA, IHUB, HQA, or HQB) and between environmental sources (sink drains versus wastewater) (**Figure 2A**).

**Figure 2.**
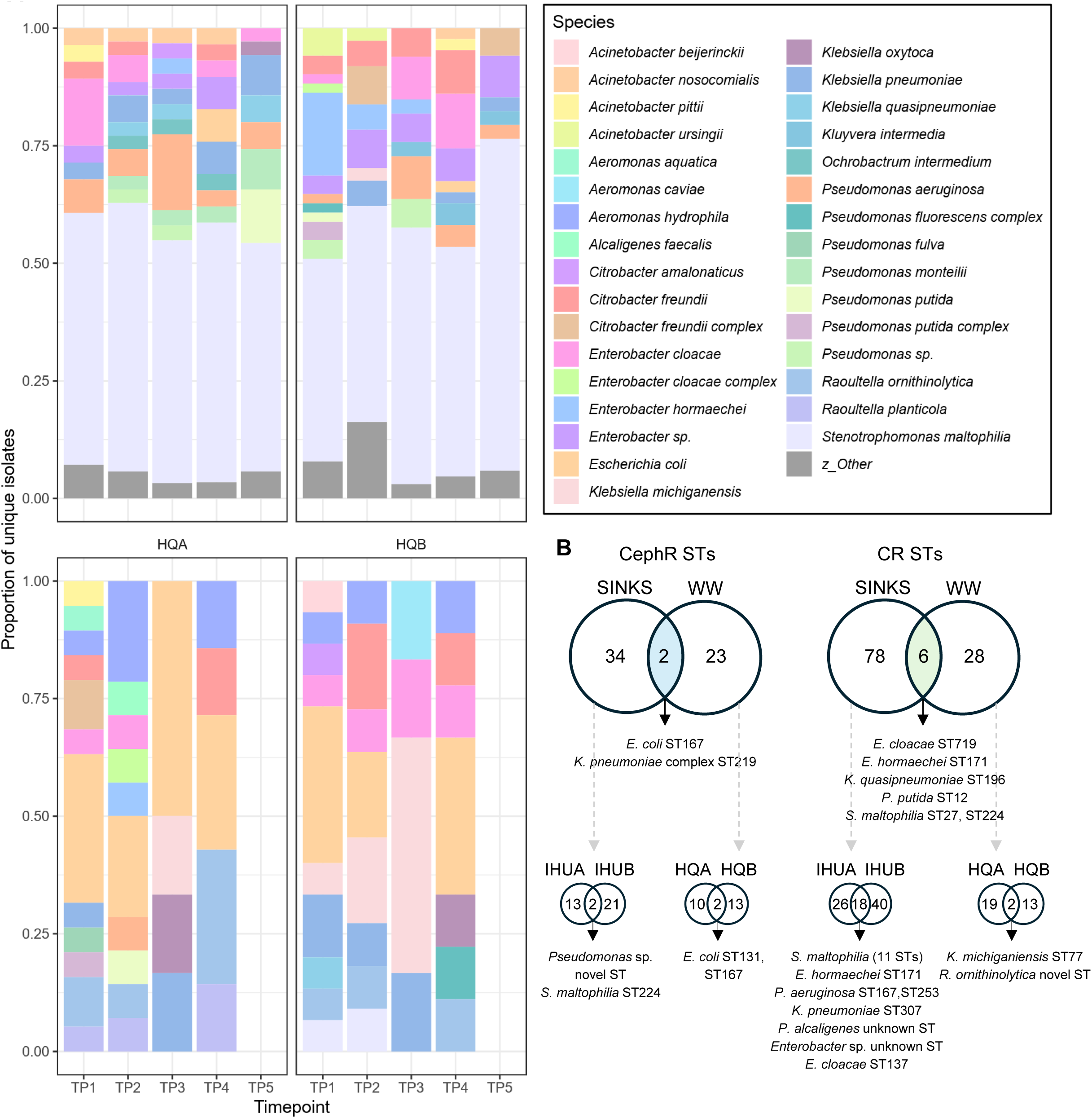
Environmental niche-specific taxonomic profiles. **(A)** Bar plots showing species diversity for each source site across sampling timepoints. For each timepoint and collection source (inpatient hospital units IHUA and IHUB, hospital wastewater quadrants HQA and HQB), the proportion of isolates assigned to a given species is displayed. All species included in the top 10 most prevalent species from at least one of the four sites are shown; all other species are grouped as “Other”. *Stenotrophomonas maltophilia* dominated sink drain multidrug-resistant gram-negative bacteria (MDR-GNB), comprising a median of 53% of total isolates (IQR: 6%) from IHUA and IHUB at a single timepoint. In contrast, wastewater-derived isolates primarily reflected common enteric species including *E. coli* and other Enterobacterales. Despite the dominance of *S. maltophilia*, sink drains routinely reflected increased species (panel A) and sequence type diversity (panel B) compared to wastewater sources. **(B)** Venn diagrams show the number of third-generation cephalosporin-resistant (CephR) or carbapenem-resistant (CR) sequence types (STs) unique to sink drains or wastewater or shared across the two source types. Niche-specific differentiation of CephR- and CR-GNB STs was evident, with limited overlap across sample source (sink drains vs. wastewater) and, within each source, collection site (IHUA vs. IHUB, HQA vs. HQB).

*Stenotrophomonas maltophilia* was the most frequently identified MDR-GNB species in sink drains, comprising a median of 53% (IQR 6%) of all isolates collected from IHUA and IHUB at a single timepoint (**Figure S4**). CR-*S. maltophilia* was the predominant phenotype (n=179 out of 288 total CR-GNB from sink drains) and was comprised primarily of ST224 (41, 23%), ST24 (19, 11%), ST229 (17, 9%) and ST233 (14, 8%) (**Figure S5**). CR-*S. maltophilia* ST224 (associated with global lineage Sm4a) was prevalent in both IHUA and IHUB (isolated from 24% of sink drains in each unit), and along with ST229 (Sm4a), and ST233 (Sm2) were also identified in the majority of timepoints (3 or more out of 5) in both inpatient units. Other lineages were localized to drains from a single unit, e.g. ST24 (Sm2), which was isolated from 22% of IHUA sinks only. Staff unit bathrooms sinks were less dominated by *Stenotrophomonas* (24% of unit bathroom isolates), and other taxa were similarly prevalent (24% *Acinetobacter*, 14% *Citrobacter*, and 14% *Enterobacter spp.*). Notwithstanding the predominance of *Stenotrophomonas* spp., each inpatient unit was characterized by a unique signature of other CR- and CephR-GNB species across time.

In contrast, wastewater-derived isolates consisted primarily of common enteric species, including CephR-*E. coli* and *Klebsiella pneumoniae* and various CRE including *Klebsiella*, *Enterobacter*, and *Raoultella* species (**Figure 2A**, **Figure S4**). The most frequently encountered CephR-GNB strain was *E. coli* ST131, which was persistent in both HQA and HQB across timepoints (**Figure S4**). Diverse *Aeromonas* and *R. ornithinolytica* STs composed a significant proportion of CR-GNB in HQA wastewater at several timepoints, while Enterobacterales (e.g., *K. michiganiensis* ST77) were predominant in HQB. Overall, wastewater samples consistently demonstrated lower species diversity than sink drain samples despite the predominance of *S. maltophilia* in the latter sample type; out of 48 unique species in all environmental isolates, 23 (48%) were found only in sink drains and 8 (17%) only in wastewater.

Pearson distance-based clustering of all recovered CephR- and CR-GNB STs at each timepoint revealed that IHUA and IHUB CephR- and CR-GNB profiles clustered together and distinctly from wastewater profiles (**Figure S5**). Furthermore, most MDR-GNB strains identified in the hospital environment were unique to a specific environmental source (sink drains or wastewater), though there were more shared CR-than CephR-GNB STs (**Figure 2B**). Of 59 unique CephR-GNB STs identified in our study, only 2 (3%) were shared between sink drains and wastewater; 34 (58%) were unique to sink drains and 23 (39%) to wastewater. Similarly, only 6 out of 112 total CR-GNB STs (5%) were shared across sources, while sink drains harbored 78 (70%) and wastewater 28 (25%) unique STs. *E. coli* ST167 and *K. pneumoniae* ST219 were the only two CephR-GNB STs common to sink drains and wastewater. Shared CR-GNB STs included *E. cloacae* complex ST171 (*E. hormaechei*) and ST719, *Klebsiella quasipneumoniae* ST196, *Pseudomonas putida* ST12, and *S. maltophilia* ST27 (Sm6) and ST224 (Sm4a).

Within sink drain or wastewater collections, environmental MDR-GNB also exhibited notable site-specific differentiation (distribution across hospital units or hospital quadrants; **Figure 2B**). Among CephR-GNB, only 2 of 36 (6%) sink drain STs were shared between IHUA and IHUB (a novel *Pseudomonas sp.* ST and *S. maltophilia* ST224) and only 2 of 25 (8%) wastewater STs were shared between HQA and HQB (*E. coli* ST131 and ST167). Sink drain CR-GNB had more overlap across sites; 18 of 84 STs (21%) were shared across inpatient units, driven largely by 11 unique *S. maltophilia* STs isolated from both units. Conversely, wastewater CR-GNB showed little concordance across hospital quadrants (2 out of 34 STs (6%)), and only *K. michiganiensis* ST77 and a novel *Raoultella ornithinolytica* ST were shared across sites.

Strain specificity for individual environmental niches was further evidenced by comparing the genomic similarity between isolates collected from the same versus different environmental locations. We compared the average nucleotide identity (ANI) for all environmental isolate pairs collected from the same sink, patient room, inpatient hospital unit, wastewater quadrant, collection site (IHUA, IHUB, HQA, or HQB), or environmental source (sink drains or hospital wastewater) versus those from different sinks, rooms, units, quadrants, sites, or sources (**Figure 3**). For each comparison, we also calculated the percent of total isolate pairs with ANI>99.5%, corresponding to the proportion of shared strains. Isolates collected from the same sink drain or room were significantly more closely related than those from different sinks or rooms (p<0.0001). 50% of isolate pairs from the same sink and 43% from the same room belonged to the same strain (pairwise ANI>99.5%), compared to 9% of isolate pairs from different sinks and 8% from different rooms. Correspondingly, based on pairwise ANI, isolates were significantly more similar when collected from the same environmental site (p<0.0001) or source (p<0.0001). This was more pronounced across environmental sources (10% of pairs from the same source with ANI>99.5% vs 4% from different sources), bolstering the evidence for niche-specific MDR-GNB colonization even within hospital water and wastewater infrastructure. These trends were largely but not universally consistent across all genera (**Figure S6**).

**Figure 3.**
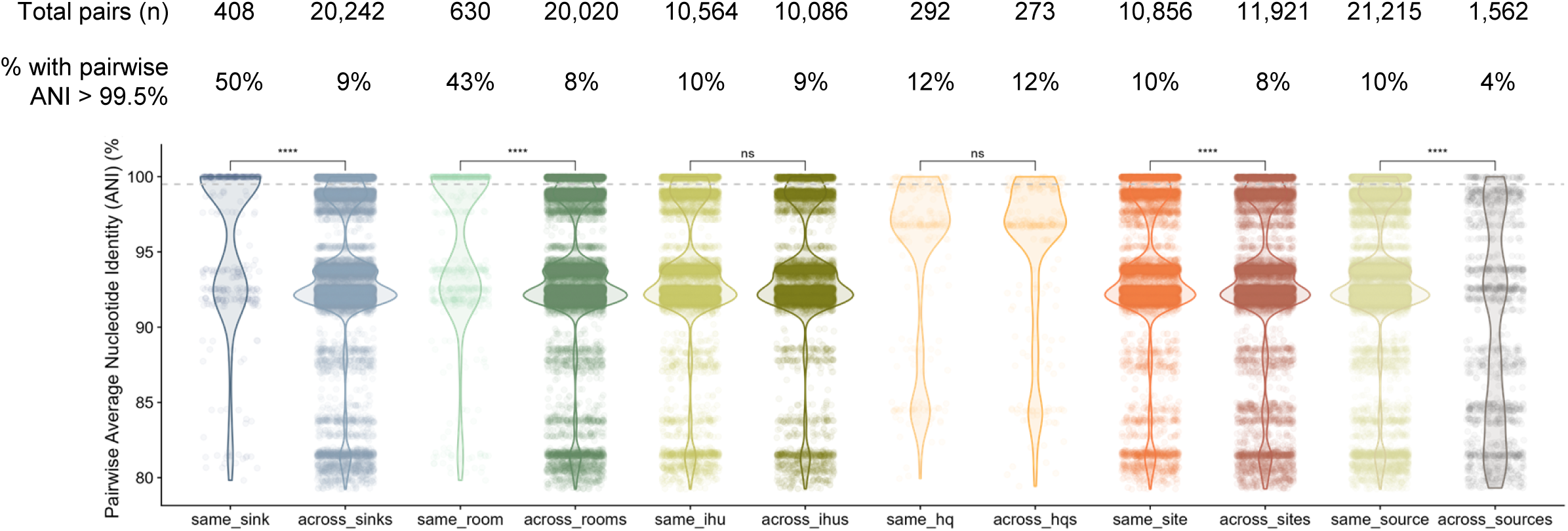
Genomic relatedness of environmental MDR-GNB isolates. Pairwise average nucleotide identity (ANI) was calculated for all sink drain and wastewater isolates belonging to the same genus. Data was then categorized based on isolate collection source and compared across categories using Wilcox tests. Isolates from the same sink (at any collection timepoint) were more closely related than sink drain isolates from different sinks (blue); to account for patient rooms with multiple sink drains, isolates collected from sink drains in the same patient room were also compared to those from different patient rooms (teal) and were more likely to be closely related. All sink drains from the same inpatient hospital unit (IHU) versus those from different IHUs (i.e., IHUA vs IHUB; olive green), as well as wastewater from the same hospital quadrant (HQ) compared to those from different HQs (i.e., HQA vs HQB; peach), did not significantly differ in terms of genomic relatedness. However, MDR-GNB from the same collection site (i.e., within IHUA, IHUB, HQA, or HQB; orange) and from the same environmental source (i.e., within sink drains or wastewater; beige) were significantly more closely related than those from different locations (maroon) or environments (grey). For each category, the total number of isolate pairs and percent of isolate pairs with pairwise ANI greater than 99.5% are also displayed; a grey dashed horizontal line indicates the ANI 99.5% threshold. (* = p<0.05; ** = p<0.01; *** = p<0.001; **** = p<0.0001)

### ARG diversity and spatiotemporal distribution

To better understand ARG spatiotemporal dynamics in hospital water systems, we analyzed the distribution of the broad-spectrum beta-lactamase genes *bla*_CTX-M_, *bla*_KPC_, and *bla*_NDM_ in environmental MDR-GNB isolates. Overall, we identified 65 *bla*_CTX-M_, 135 *bla*_KPC_, and 8 *bla*_NDM_ genes in 56, 96, and 8 environmental isolates, respectively. The most common *bla*_CTX-M_ allele was *bla*_CTX-M-15_ (47/65, 72%), while other variants were less common (*bla*_CTX-M-55_: 7/65, 11%; *bla*_CTX-M-27_: 5/65, 8%). Similarly, *bla*_KPC-3_ was the predominant *bla*_KPC_ allele (100/135, 74%), followed by *bla*_KPC-2_ (26/135, 19%) and *bla*_KPC-4_ (5/135, 4%); *bla*_NDM-1_ was the only *bla*_NDM_ variant detected. Consistently, a higher proportion of wastewater MDR-GNB were positive for *bla*_CTX-M_, *bla*_KPC_, and *bla*_NDM_ compared to those from sink drains. Out of 68 CephR isolates from sink drains and 41 from wastewater (**Table 1**), 27 (40%) and 29 (71%) were *bla*_CTX-M_ positive, respectively. Similarly, 51/288 (18%) CR sink drain and 45/46 (98%) CR wastewater isolates harbored *bla*_KPC_, and 2 (0.7%) sink drain and 6 (13%) wastewater CR isolates were positive for *bla*_NDM_.

Overall, the predominant bacterial species harboring beta-lactamase or carbapenemase genes differed across environmental sampling sources. In sink drain samples, *bla*_CTX-M_ was most commonly identified in *Enterobacter spp.* (11/27, 41%), particularly *E. hormaechei* ST134 (n=8) followed by *K. pneumoniae* (12/27, 44%). The majority of wastewater isolates harboring *bla*_CTX-M_ belonged to *E. coli* (22/29, 76%), including *E. coli* ST131 (n=8). Similarly, *bla*_KPC_ found in sink drains occurred exclusively in Enterobacterales, of which the most common were diverse *Enterobacter* sp. (33/51 isolates, including 16 *E. cloacae* and 5 *E. hormaechei*) followed by *Citrobacter* sp. (12/51 isolates). Wastewater isolates harboring *bla*_KPC_ (n=45) primarily included isolates belonging to opportunistic *Aeromonas* sp. (n=10) and *Pseudomonas aeruginosa* (n=1); the most frequently identified *bla*_KPC_-harboring Enterobacterales belonged to the *Raoultella* genus (n=11/45, including 8 *R. ornithinolytica* and 3 *R. planticola*). *bla*_NDM_ was found in *Citrobacter sp. a*nd *K. oxytoca* isolates in both environmental sources, as well as *K. pneumoniae*, *Enterobacter sp.,* and *Raoultella sp.* in wastewater.

When we examined specific sampling locations over time, we found that *bla*_CTX-M-15_ was identified in both hospital quadrants across all four wastewater sampling timepoints and persisted in three different hospital room sink drains (**Figure 4**). No other *bla*_CTX-M_ variants showed evidence of persistent environmental colonization. Similarly, *bla*_KPC-3_ was detected in all wastewater samples, while *bla*_KPC-2_ was detected across all HQB and 3/4 HQA sampling timepoints. *bla*_KPC-3_ was also persistently detected in 9 hospital room sink drains. *bla*_NDM-1_ detection in wastewater was limited to samples from HQA but was identified across sampling timepoints; it was also identified in sink drains from both hospital units (IHUA and IHUB) at timepoint 5.

**Figure 4.**
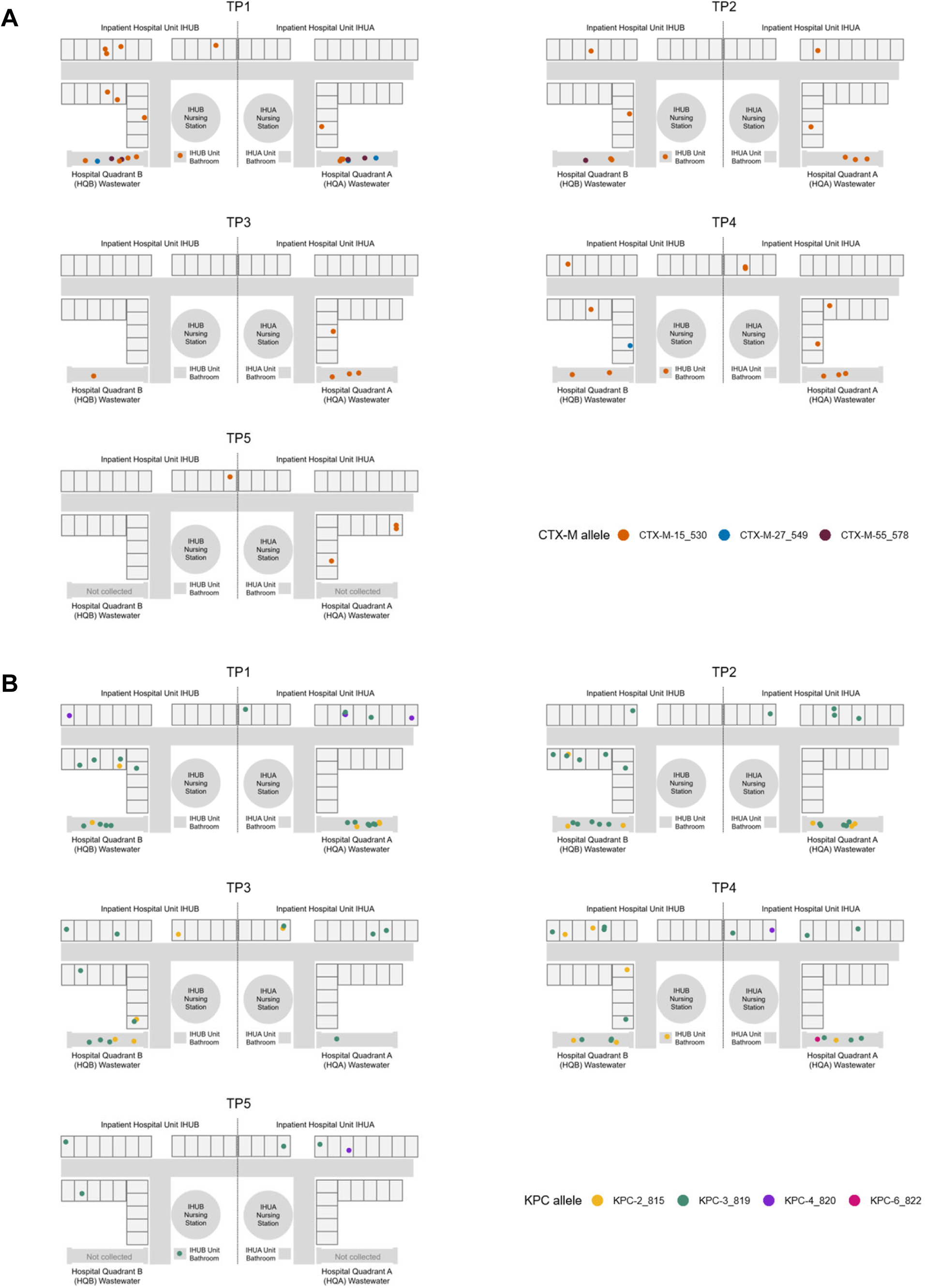
Spatiotemporal analysis of ARG allele distribution. To assess the spatial distribution of resistance gene alleles across timepoints, (**A**) *bla*_CTX-M_ and (**B**) *bla*_KPC_ alleles identified in environmental isolates were plotted based on isolate collection location. Persistence within a sink drain was defined based on identification in the same sink drain over at least two collection timepoints. (**A**) Of the three *bla*_CTX-M_ alleles in our collection, *bla*_CTX-M-15_ was the most prevalent and was identified in both hospital quadrants across all wastewater sampling timepoints (4/4 timepoints). *bla*_CTX-M-15_ also persisted in three different hospital room sink drains. Other variants, *bla*_CTX-M-27_ and *bla*_CTX-M-55_, were almost exclusively associated with wastewater samples and did not show evidence of persistent environmental colonization. (**B**) The most prevalent *bla*_KPC_ alleles in our collection, *bla*_KPC-3_ and *bla*_KPC-2_, were detected in nearly all wastewater samples (*bla*_KPC-3_ in all wastewater samples; *bla*_KPC-2_ in 3/4 HQA sampling timepoints and 4/4 HQB timepoints). *bla*_KPC-3_ was also persistently detected in 9 hospital room sink drains.

Although we focused our analyses on *bla*_CTX-M_, *bla*_KPC_, and *bla*_NDM_ as the primary drivers of cephalosporin or carbapenem resistance in our environmental collection, other broad-spectrum beta-lactamase genes were also detected in environmental isolates. In addition to *bla*_NDM_, *bla*_VIM_-type metallo-beta-lactamases were identified in 11 *Pseudomonas* sp. sink drain isolates, of which 8 were *bla*_VIM-2_–harboring *P. aeruginosa* ST253 distributed across HUA rooms and sampling timepoints. *bla*_VIM-4_ and *bla*_IMP-13_ were detected in wastewater samples in *Alcaligenes faecalis* and *A. hydrophila,* respectively. Diverse Enterobacterales harboring non *bla*_CTX-M_-ESBL genes included 22 environmental isolates harboring variants of *bla*_SHV_ with ESBL activity.^27^ Of these, *bla*_SHV-30_ was the most common, and was found in multiple Enterobacterales species including *C. freundii,* ST166, *R. planticola, Kluyvera intermedia,* and *Enterobacter* sp. MDR-GNB from both sink drain and wastewater isolates also harbored acquired ARGs conferring resistance to a variety of other drug classes, including aminoglycosides, fluoroquinolones, and sulfonamides (data not shown).

### Diverse ARG-harboring plasmid clusters in environmental sources

As expected, our ARGs of interest were overwhelmingly plasmid-borne. Out of 65 *bla*_CTX-M_ genes, 39 (60%) were located on plasmids, including 18 plasmids from sink drains and 21 from wastewater. 102 of 135 (76%) copies of *bla*_KPC_ were identified on 97 plasmids, of which 50 were isolated from sink drains and 47 from wastewater. *bla*_NDM_ (n=8) was exclusively located on 2 plasmids isolated from sink drains and 6 from wastewater.

To further characterize ARG-harboring plasmids, we extracted plasmid contigs for clustering analysis based on pairwise genetic distances and mobility prediction. *bla*_CTX-M_-harboring plasmids formed 10 primary clusters (applying Mash distance-based clustering with cutoff of 0.06; 1-16 plasmids per cluster) and 15 secondary clusters (cutoff of 0.025; 1-9 plasmids per cluster) (**Figure S7A**). The majority of *bla*_CTX-M_-harboring plasmids (11/15 secondary clusters comprising 29/39 plasmids) belonged to the IncF plasmid family. However, a cluster of 5 IncHI2A plasmids encoding *bla*_CTX-M-15_ were isolated from sink drains, and sporadic IncC, IncI-γ/K1, and nontypeable plasmids were also identified. *bla*_KPC_ was encoded by diverse plasmids belonging to 29 primary and 41 secondary clusters (1-16 plasmids per primary cluster; 1-14 plasmids per secondary cluster), including 25 singletons based on secondary cluster designations. IncF-type plasmids comprised the largest number of plasmid clusters (19 clusters including 32 plasmids) followed by nontypeable (11 clusters with 29 plasmids) and IncN plasmids (2 clusters with 15 plasmids), as well as smaller clusters of IncC, IncP, IncX5, and ColRNAI plasmids. *bla*_KPC_ was also found on 9 co-integrated plasmids with a variety of *rep* gene types, the majority of which were found in wastewater isolates (7/9, 78%), suggesting increased plasmid recombination in this setting. *bla*_NDM_-harboring plasmids formed 2 primary and 3 secondary clusters; the three secondary clusters included IncX3 (n=4/8) and IncU-type plasmids (n=3/8), and a nontypeable singleton. The majority of plasmids encoding *bla*_CTX-M_ (35/39, 90%), *bla*_KPC_ (68/97, 67%), and *bla*_NDM_ (100%) were predicted to be conjugative or mobilizable (**Figure S7B**).

Among *bla*_CTX-M_ plasmids, only one primary (10%) and 2 secondary clusters (13%) included plasmids found in multiple species (**Figure S7A**). CTX-M AA028 (IncFIB/FII), was identified in *K. pneumoniae* and *K. michiganensis* isolates, while CTX-M AA027 (IncFII) was the only secondary cluster consisting of plasmids found in multiple genera (*K. pneumoniae*, *R. planticola*, and *E. coli)*. In contrast, 12 *bla*_KPC_ primary clusters (41%) and 7 secondary clusters (17%) comprised plasmids from at least two different species. Of these, KPC AA066 (nontypeable) and KPC AA053 (IncN) were comprised of isolates from 6 and 4 Enterobacterales species, respectively. Both primary clusters and 2 of 3 secondary clusters of *bla*_NDM_ plasmids contained plasmids found in three distinct Enterobacterales species.

Although wastewater and sink drains harbored primarily distinct bacterial strains, plasmid sharing across environmental sources occurred frequently; 3/15 (20%) *bla*_CTX-M_-harboring secondary plasmid clusters, 6/41 (15%) *bla*_KPC_ secondary clusters, and 1/3 (33%) *bla*_NDM_ secondary clusters included plasmids found in both wastewater and sink drain samples (**Figure S8**). When we examined individual plasmid clusters across timepoints, we found evidence for gene and source-specific differences in prevalence and persistence (**Figure S9**). Out of 15 *bla*_CTX-M_ plasmid secondary clusters, 4 (27%) were identified over at least two timepoints in the same environmental source (wastewater or sink drains), only one of which (AA028) was both persistent in IHUA and IHUB and found in wastewater and sink drains (**Figure S10**). 14 out of 41 (34%) *bla*_KPC_ and 2 out of 3 (67%) *bla*_NDM_ secondary clusters also persisted over time. However, only five *bla*_KPC_ secondary clusters, KPC AA053, AA065 (nontypeable), AA066, AA067 (IncX5), and AA078 (IncC), were persistent in both sink drains and wastewater. In several cases, we observed contemporaneous overlap of the same *bla*_KPC_ plasmid secondary cluster in sink drains and wastewater connected by hospital plumbing lines; for example, KPC AA067 was identified in the IHUB unit and only HQB wastewater (**Figure S10**). Despite consistent growth of MDR-GNB from both inpatient unit staff bathrooms across all timepoints, no *bla*_CTX-M_ plasmids and just two *bla*_KPC_ plasmids (KPC AA066 and AA089 harbored by *Citrobacter* spp.) were isolated from the unit bathroom sink drain in IHUB, and no ARG-harboring plasmids were recovered from the unit bathroom in IHUA.

### Relationships between environmental and clinical MDR-GNB and antibiotic resistance plasmids

In our previous work, close phylogenetic clustering of clinical isolates collected years apart from our hospital center, without any clear epidemiological links, indicated the likelihood of unsampled reservoirs.^20^ Therefore, to assess long-term persistence of clinically relevant MDR-GNB in the hospital environment, we compared the relatedness of isolates between environmental samples from this study and a large, longitudinal collection of 602 historical MDR Enterobacterales clinical isolates from the same hospital between 2009-2020. Across these collections, we identified 18 genera, including 13 only found in environmental sources and five Enterobacterales genera containing both environmental and clinical isolates. For all isolate pairs within each genus, we used pairwise ANI to define genomic clusters (ANI>99.5%), which we then quantified and classified by collection source(s) (**Figure 5A**). Only four genera (22%), *Enterobacter, Klebsiella, Pseudomonas,* and *Stenotrophomonas*, included genomic clusters with isolates from both sinks and wastewater, reflecting limited evidence for an MDR-GNB signature specific to hospital plumbing. Genomic network analysis of the five Enterobacterales genera with overlap across environmental and clinical collections revealed multiple instances of strain sharing across collection source (**Figure 5B**). Within each genus, 1-8 clusters (10-22% of total clusters per genus) contained both environmental (sink drain, wastewater, or both) and clinical isolates, and these clusters included a significant proportion of Enterobacterales isolates (225/602 (37%) of total Enterobacterales). Three Enterobacterales genera (*Enterobacter*, *Escherichia*, and *Klebsiella*) each formed one genomic cluster consisting of isolates from all three sources (sink drains, wastewater, and clinical).

**Figure 5.**
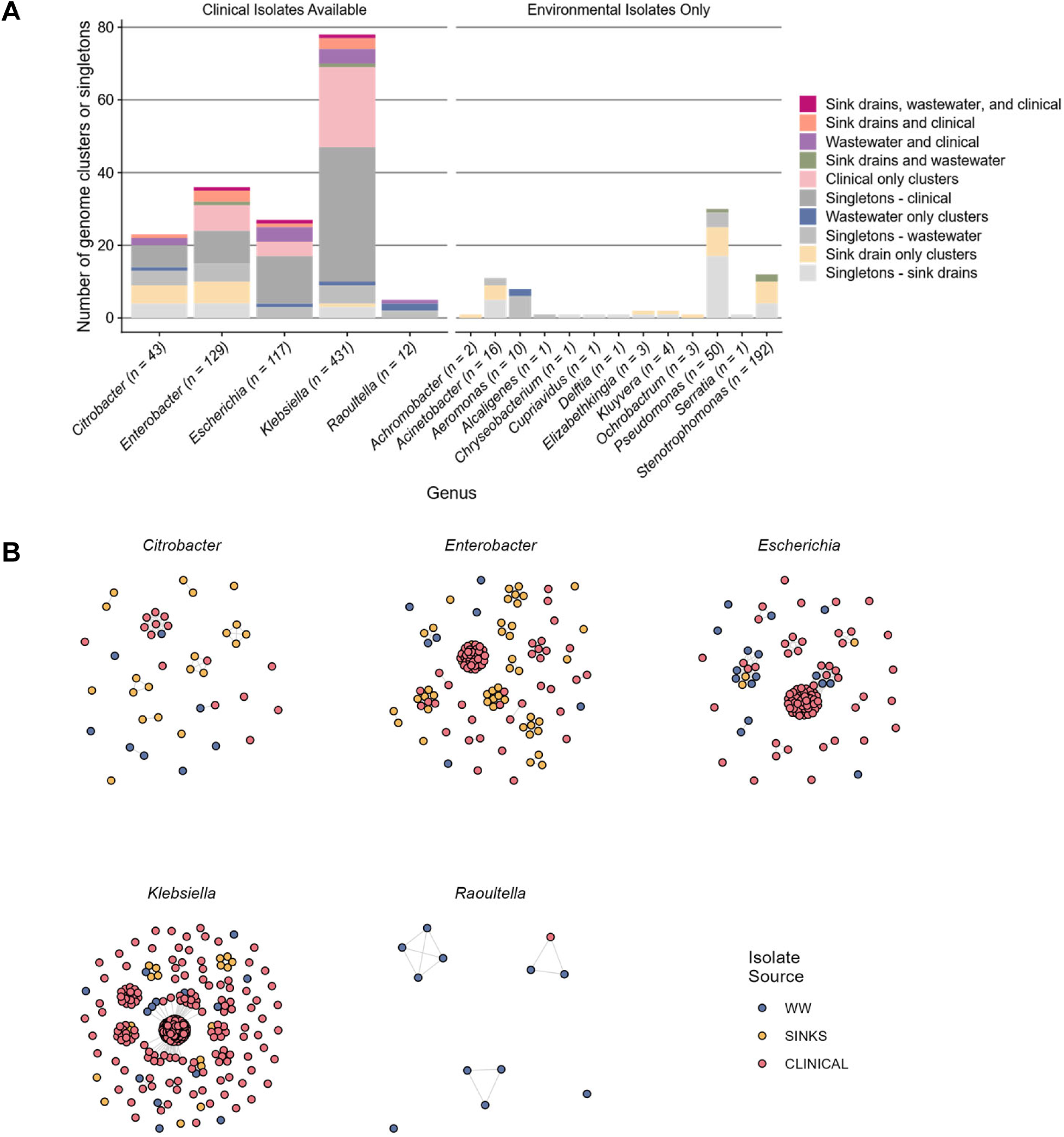
Genomic relatedness of environmental and clinical MDR-GNB. For each genus in our environmental dataset, we performed network analyses based on pairwise average nucleotide identity (ANI) to identify closely related genomic clusters and singletons (isolates with no other closely related neighbors). Pairwise ANI of 99.5% was used as a threshold to define genomic clusters. (**A**) Stacked bar plots show the sources of all genomic clusters and singletons. The height of each bar indicates the total number of genomic clusters and singletons identified for the genus. Bottom axis labels indicate the genus and the number of isolates included in the network analysis. Colors are used to differentiate the iMost genomic clusters were singletons or derived from a single source (100% of genomic clusters in 11 of 18 genera (61%)). Genomic clusters shared between sink drains and wastewater were rare (4 of 18 genera (22%)), reflecting limited evidence for a specific hospital plumbing signature of MDR-GNB. For five genera, as indicated in the labels above the bar plots, both environmental and clinical MDR-GNB were available for analysis, while the remaining 13 genera comprised of environmental isolates only. The five Enterobacterales genera with clinical data universally showed evidence solate source(s) for each cluster or singleton. of strain sharing between environmental and clinical collections (10-22% of total clusters within each genus). (**B**) For the five Enterobacterales genera identified in both clinical and environmental MDR-GNB collections, network plots were generated to visualize relationships between sink drain, wastewater, and clinical genomes; each point represents an assembled bacterial chromosome, and colors indicate isolate collection source. Multiple examples of environmental genome clusters, with isolates from sink drains and wastewater, and environmental-clinical clusters, with isolates from clinical and at least one environmental source, were identified.

Re-clustering of plasmid sequences to also include both environmental and clinical ARG-harboring plasmids resulted in the identification of 51 *bla*_CTX-M_ plasmid secondary clusters (including 25 singletons), 85 secondary clusters of *bla*_KPC_ plasmids (38 singletons), and 4 secondary clusters of *bla*_NDM_ plasmids (1 singleton); 3 plasmids encoding both *bla*_CTX-M_ and *bla*_NDM_ comprised overlapping *bla*_CTX-M_ and *bla*_NDM_ plasmid clusters (CTX-M BB098 and NDM BB005). Among 140 total ARG-harboring plasmid clusters, 25 (18%) were shared between at least two sources, and 5 (4%) were shared across all three sources (**Figure 6**). Not surprisingly, clusters present in all three sources included several of the largest plasmid clusters, such as KPC BB010 (IncN), which consisted of 38 plasmids and CTX-M BB050 (IncFIB/FII) with 23 plasmids.

**Figure 6.**
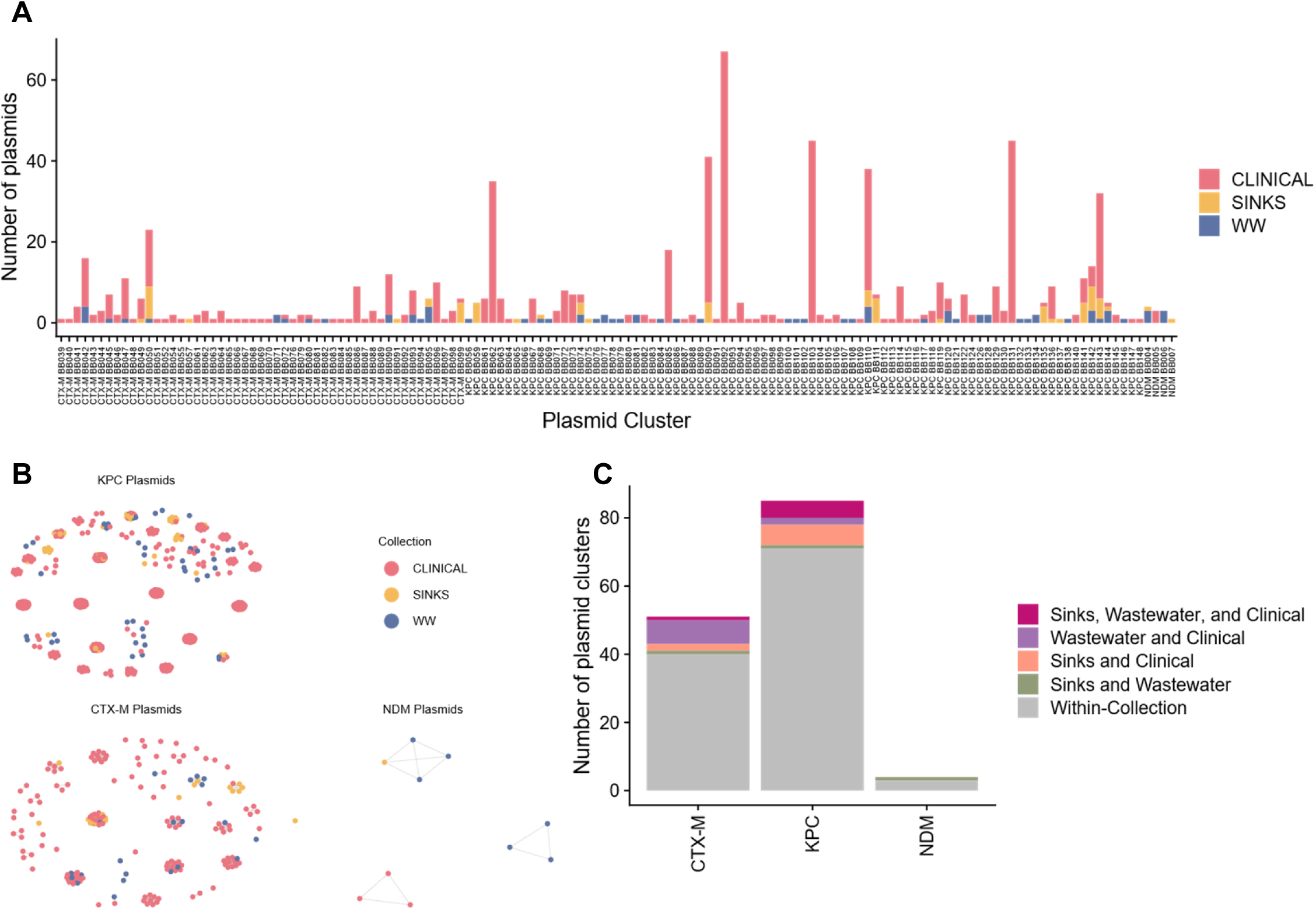
Environmental and clinical antibiotic resistance plasmid clusters. Plasmids harboring *bla*_CTX-M_, *bla*_KPC_, or *bla*_NDM_ from environmental and clinical collections were grouped into secondary clusters by MOB cluster (Mash distance <=0.025). (A) For each cluster, bar plots indicate the total number of plasmids from each collection source (sink drains, wastewater (WW), or clinical; indicated by color). (B) Network plots based on secondary cluster membership show each plasmid as a single dot, with color indicating collection source. (C) For each of the three antibiotic resistance genes, bar plots summarize the number of plasmid secondary clusters which were identified in only one source (within-collection), shared across environmental sources (sink drains and wastewater), or shared between environmental and clinical collections (sink drains and/or wastewater and clinical). 10/51 (20%) of *bla*_CTX-M_ and 13/85 (15%) of *bla*_KPC_ plasmid clusters were shared between at least one environmental source and clinical isolates.

Among plasmid clusters including plasmids from two different sources, 22 consisted of plasmids shared between clinical and wastewater or sink drain isolates, whereas only 8 were shared between environmental sampling sources. Interestingly, a greater proportion of *bla*_CTX-M_ plasmid clusters were shared between wastewater and clinical isolates (8/51, 16%) than between sink drains and clinical isolates (3/51, 6%), while the reverse was true for *bla*_KPC_ plasmids (7/85 (8%) shared between clinical and wastewater isolates vs. 11/85 (13%) between clinical and sink drain isolates). Most plasmid clusters, regardless of ARG, were isolated from a single source (40/51 (78%) for *bla*_CTX-M_, 71/85 (84%) for *bla*_KPC_, and 3/4 (75%) for *bla*_NDM_). Reframed by environmental source, 13/21 (62%) *bla*_CTX-M_ and 17/47 (36%) *bla*_KPC_ plasmids from wastewater, versus 14/18 (78%) *bla*_CTX-M_ and 41/50 (82%) of *bla*_KPC_ plasmids from sinks, clustered with plasmids from clinical isolates.

### Quantifying MDR-GNB and antibiotic resistance plasmids links across environmental reservoirs and clinical infections

Finally, we aimed to comprehensively quantify links between MDR-GNB strains and ARG-harboring plasmids across environmental and clinical collections (**Figure 7**). First, we calculated the proportion of isolates from each inpatient unit or wastewater quadrant that shared a chromosome, plasmid, or both across collection sites. While a high proportion of isolates shared a chromosome across environmental sites (n=237/443, 53%; 107/158 (68%) IHUA, 95/198 (48%) IHUB, 14/46 (30%) HQA, and 21/41 (51%) HQB isolates), this was largely driven by genome sharing within the same environmental source (between sink drains across inpatient units or between hospital wastewater quadrants), consistent with other analyses in this study. Isolates sharing both a chromosome and plasmid across environmental sources were rare (n=14/443, 3%), comprising only 6 (2%) sink drain and 8 (9%) wastewater isolates.

**Figure 7.**
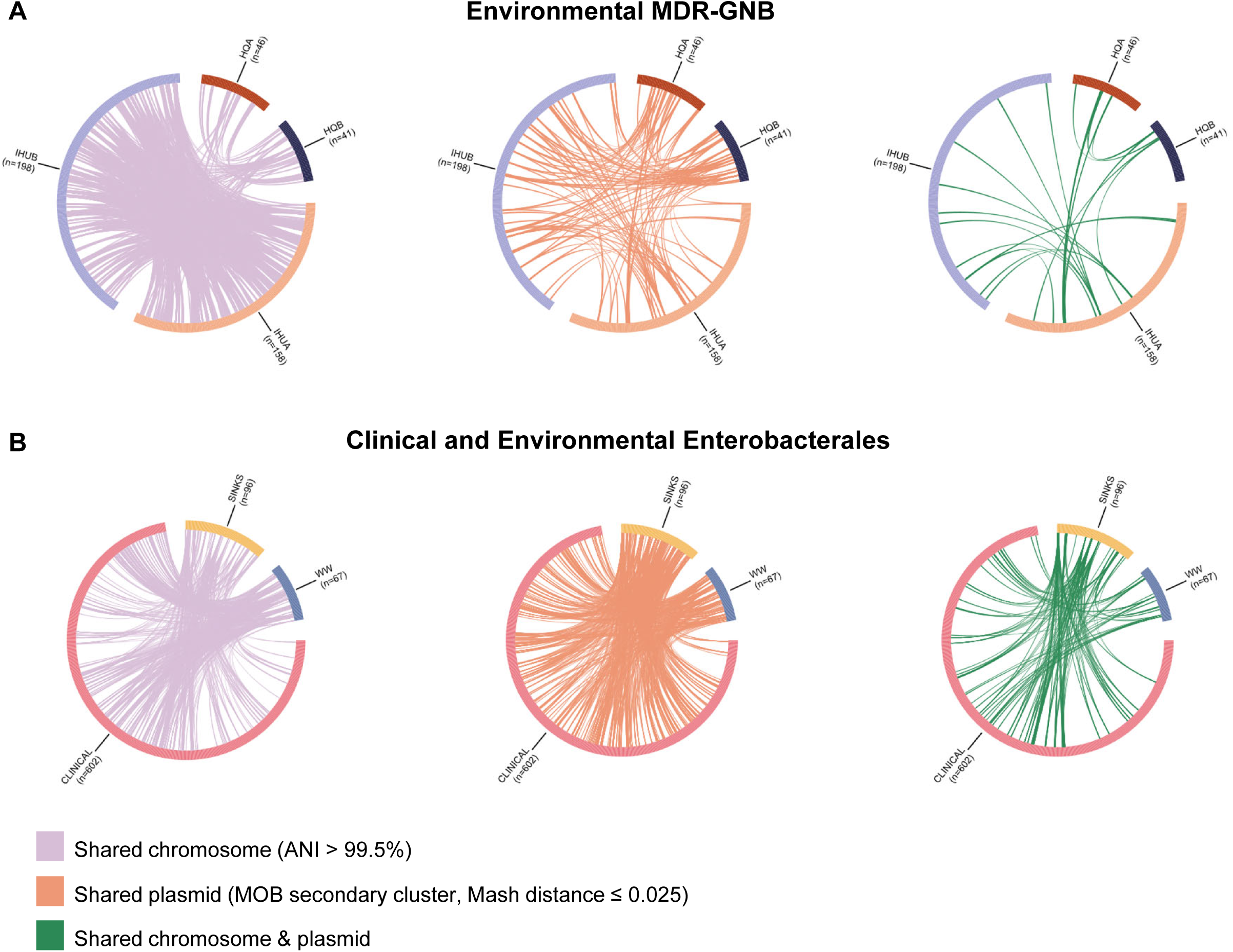
MDR-GNB strain and antibiotic resistance plasmid sharing between environmental reservoirs and clinical isolates. All pairs of isolates within our environmental collection (A) or all pairs of Enterobacterales isolates from both clinical and environmental collections (B) were assessed to determine whether the pair shared a chromosome (based on pairwise ANI) or plasmid (based on MOB secondary cluster assignment) only, or both. Outer rings of each circle plot represent all isolates from a given environmental collection site (panel A; IHUA, IHUB, HQA, or HQB) or collection source (panel B; sink drains (SINKS), wastewater (WW), or clinical). Leftmost plots show lilac lines connecting each isolate pair with only shared chromosomes (chromosome pairwise ANI > 99.5% but plasmid pairwise Mash distance > 0.025). Middle plots show peach lines connecting each isolate pair with differing chromosomes but harboring the same plasmid (same plasmid secondary cluster with Mash distance ≤ 0.025 but chromosome pairwise ANI ≤ 99.5%). Rightmost plots show green lines connecting each isolate pair sharing both a chromosome and plasmid (chromosome pairwise ANI > 99.5% and same plasmid secondary cluster); Only links across environmental collection sites (A) or between sources (B) are shown; links from the same site or source are not displayed.

Our data further supported the differing roles of chromosome versus plasmid sharing between environmental sites when analyzed in terms of related isolate pairs (**Figure 7A**). Given 443 unique environmental isolates included in this study, there were a total of 97,903 unique isolate pairs, including 64,142 pairs collected from different environmental sites (IHUA, IHUB, HQA, or HQB). Of these, 2,037 isolate pairs (2.1% of all unique pairs) shared a chromosome, driven primarily by *S. maltophilia* found in sink drains, and only 947 isolate pairs sharing a chromosome were isolated from different collection sites (1.5% of all pairs from different sites). 277 (0.3%) environmental isolate pairs shared an ARG-harboring plasmid, including 147 (0.2%) collected from different environmental sites, largely due to plasmid sharing by Enterobacterales strains. Lastly, 97 (0.2%) pairs shared both a chromosome and an ARG plasmid, including 27 (0.04%) pairs from different sites. When we reanalyzed isolate pair data considering only Enterobacterales species (n=163), we found that of 9,662 total environmental Enterobacterales isolate pairs across sites, 73 (0.8%) shared a chromosome, 144 (1.5%) shared a plasmid, and 27 (0.3%) shared both. Few pairs consisting of a sink drain and a wastewater Enterobacterales isolate shared a chromosome (23 of 6,432 total pairs; 0.4%), plasmid (92; 1.4%), or both (10; 0.2%).

Next, we explored connections between environmental and clinical Enterobacterales by calculating the proportion of wastewater (n=67) or sink drain (n=96) isolates which shared a chromosome, plasmid, or both with at least one clinical isolate (n=602). Overall, 32 (33%) sink drain and 30 (44%) wastewater Enterobacterales isolates shared chromosomes with clinical isolates, and 55 (57%) sink drain and 28 (42%) wastewater isolates harbored clinically associated plasmids. 14 (15%) sink drain and 6 (9%) wastewater Enterobacterales shared both a chromosome and plasmid with at least one clinical isolate. A higher proportion of clinical isolates shared strains (chromosomes) with wastewater (131; 22%) compared to sink drains (76; 13%), including 44 (7%) clinical isolates linked with both environmental sources. 144 (24%) clinical isolates had plasmids that clustered closely with plasmids found in sink drains and 128 (21%) shared plasmids with wastewater isolates. Significant proportions of clinical plasmids harboring *bla*_CTX-M_ (51/137; 37%) and *bla*_KPC_ (123/447; 28%) were also identified in environmental sources. Clinical isolates that shared both a chromosome and plasmid with isolates from sink drains (42; 7% of all clinical isolates) and wastewater (18; 3%) largely included epidemic CRE and ESBLE clones (e.g. *E. hormaechei* ST171, and *K. pneumoniae* ST307 in clinical and sink drain isolates; *E. coli* ST131 and *K. pneumoniae* ST17 in clinical and wastewater isolates). Plasmids from clinical isolates found in both sink drains and wastewater primarily belonging to secondary clusters KPC BB110 (IncN; 30/78 isolates, 38%), KPC BB143 (nontypeable; 26/78, 33%), and CTX-M BB050 (IncFIB/FII; 14/78, 18%).

Fewer invasive clinical isolates collected from blood, abdominal, intraabdominal, or respiratory sources shared a chromosome with environmental isolates compared to those from non-invasive sources including rectal swabs, urine, and wounds (24% vs 30%); however, the proportion of invasive isolates with linked genomes to environmental sources steadily increased over time (5% in 2011 to 35% in 2018), while the proportions for non-invasive isolates remained stable over time (median 26%) (**Figure S11**). Plasmid sharing between clinical and environmental isolates, in contrast, was similar for invasive (33%) and non-invasive (32%) clinical sources, although plasmid sharing by invasive isolates also increased over time (17% to 55%, 2010-2018).

Lastly, we analyzed environmental and clinical Enterobacterales isolate pairs to quantify the rates of chromosome and plasmid sharing between environmental and clinical sources (**Figure 7B**). Again, chromosome and plasmid links between environmental and clinical MDR-GNB were more prevalent than those between isolates collected from different environmental sources. A total of 765 Enterobacterales isolates (163 environmental and 602 clinical isolates) comprised 292,230 unique isolate pairs including 104,558 pairs between sink drain, wastewater, or clinical sources. Chromosome sharing was detected in 33,643 (11.5%) Enterobacterales isolate pairs, including 872 pairs between environmental and/or clinical sources (0.8% of all pairs across sources). Similarly, 8,152 (2.8%) of Enterobacterales pairs harbored plasmids belonging to the same resistance plasmid cluster, including 1,000 (1.0%) across collections. 4,515 (1.6%) pairs shared both a chromosome and plasmid, including 212 (0.2%) between hospital environment and clinical sources. Of 40,334 Enterobacterales wastewater-clinical isolate pairs, 557 (1.4%) shared a chromosome, 287 (0.7%) shared a plasmid, and 33 (0.08%) shared both, while 292 of 57,792 (0.5%) sink drain-clinical isolate pairs shared a chromosome, 646 (1.1%) shared a plasmid, and 169 (0.2%) shared both. There was no significant difference in the proportion of sink drain versus wastewater isolates sharing a chromosome with clinical isolates (*p*=0.2). A higher proportion of sink drain isolates compared to wastewater isolates demonstrated plasmid sharing with clinical isolates, but the difference did not reach statistical significance (*p*=0.07).

## Discussion

In this study, we examined the population, spatial, and temporal dynamics driving the abundance and persistence of MDR-GNB across hospital water systems, critical reservoirs for MDR bacteria in healthcare systems. We detected widespread contamination of hospital sinks and wastewater streams with CR- and CephR-GNB, consisting of both intrinsically beta-lactam resistant environmental bacteria (e.g. *Stenotrophomonas* sp.) and predominantly enteric bacteria harboring ARGs (e.g. *bla*_KPC_ and *bla*_CTX-M_). Within each site, the distribution and persistence of ARGs was associated with specific bacterial subpopulations, supporting an important role for niche-specific factors in facilitating colonization. Moreover, we identified links between environmental and historical clinical isolates that were similarly niche- and ARG-specific, further emphasizing differences across sources. Finally, by applying comprehensive nanopore sequencing, we found evidence for transmission of resistance plasmids across sources, connecting bacterial reservoirs and demonstrating how the interchange of ARGs amplifies the burden of healthcare-associated MDR-GNB. Taken together, our findings highlight the unique benefits of complementary sink drain and wastewater surveillance and the need for complete plasmid sequencing to fully characterize environmental reservoirs for MDR-GNB.

Rather than limiting our analysis to specific resistance categories or organisms (e.g. CRE), we included all culturable MDR-GNB, facilitating characterization of diverse bacteria within each environmental source. In sink drains, MDR-GNB colonization was dominated by opportunistic bacteria such *S. maltophilia* and *P. aeruginosa* that harbor a variety of intrinsic and acquired mechanisms of antibiotic resistance and are well-adapted to persist in this environment.^8,33^ Widespread colonization of sink drains with *S. maltophilia* which was linked in prior studies to persistence of distinct genotypes associated with biofilm formation and nutrient utilization.^34^ Specific Enterobacterales lineages such as *E. cloacae* complex and *Citrobacter* sp. were prevalent in sink drains and, concerningly, frequently harbored *bla*_CTX-M,_ *bla*_KPC_, and *bla*_NDM_, suggesting that they may be important reservoirs for ARGs in this setting. Common enteric bacteria such as *E. coli* and *K. pneumoniae* were infrequently detected in sink drain samples but accounted for most MDR-GNB in wastewater, where they were also associated with broad-spectrum beta-lactamase gene carriage. While opportunistic bacteria harboring ARGs were relatively rare, we detected *bla*_KPC_ in diverse *Aeromonas* sp. including one isolate that shared a *bla*_KPC –_harboring plasmid with other Enterobacterales, suggesting a role for occult ARG carriage in select environmental bacteria. Niche-specific differentiation of CephR- and CR-GNB STs was also evident, with limited overlap across sample source (sink drains vs. wastewater) and, within each source, collection site (IHUA vs. IHUB, HQA vs. HQB). Differences across these microenvironments may have been due to ward- and/or quadrant-specific differences in patient comorbidities, selection pressure, and plumbing architecture. Overall, these findings suggest that hospital plumbing infrastructure consists of multiple linked components, each with a distinct microbiome, rather than representing a single reservoir.

Using a high-throughput nanopore sequencing pipeline for MDR-GNB isolates cultured from environmental sources, we were able fully characterize shared *bla*_CTX-M,_ *bla*_KPC_, and *bla*_NDM_-harboring plasmids. Based on comparative analysis with historical clinical isolates, we also assessed long-term persistence of plasmids previously associated with clinical infections. Overall, plasmid sharing across environmental sources accounted for a greater proportion of linked isolate pairs than chromosome sharing, although individual plasmid clusters were largely derived from a single environmental source rather than found in both sink drain and wastewater isolates. However, there were plasmid clusters that demonstrated remarkable transmissibility across environmental sources and bacterial organisms, most notably the *bla*_KPC_-harboring IncN plasmid BB010, which we previously found to be highly promiscuous and conserved within our hospital system.^35^ We also found evidence that plasmids can facilitate the spread of ARGs between distinct environmental niches, including via transmission to well-adapted environmental organisms such as *Aeromonas, Citrobacter, Enterobacter, and Raoultella* spp., which has been documented in prior studies.^22^ Finally, we found evidence for remarkable persistence of resistance plasmids for years following initial detection in clinical isolates. This highlights the need for both chromosome and plasmid surveillance to better understand the prevalence of pathogenic strains versus the risk of diversification into diverse bacterial hosts for a given resistance plasmid.

This study had many strengths, in particular our longitudinal sink drain and wastewater sampling from corresponding hospital quadrants to enable comparison of diverse MDR-GNB across sites and sources and over time. We also comprehensively applied nanopore sequencing to assess MDR-GNB and resistance plasmids across multiple sample types. We also acknowledge several limitations. All colonies from selective growth media were not sequenced comprehensively; instead, to optimize for MDR-GNB diversity, we sequenced select colonies from each unique morphology observed. Thus, we did not systematically investigate genetic diversity within individual organisms from the same environmental sample. We also did not assess patient colonization, as the central aim of this study was to compare the population structure of MDR-GNB within environmental niches. Clinical and environmental isolates were not contemporaneous, as clinical isolates analyzed in this study predated those from the hospital environment by 6-17 years. However, our prior data show the potential for clinical strains to reemerge after many years^14,20^ and current findings further support the relevance of historical strains and resistance plasmids to contemporary hospital-associated MDR-GNB populations. Finally, we did not assess nonculturable bacteria and organisms lacking specific antibiotic-resistance phenotypes, e.g. using metagenomic sequencing, which may contribute to the persistence of MDR-GNB by serving as occult reservoirs or ARGs or forming favorable microbial communities.

In summary, the findings of our study underscore differences in the prevalence and composition of MDR-GNB within distinct components of the hospital built environment and their potential to contribute to MDR-GNB transmission and persistence. Thus, while this study supports the role of the hospital environment as an important reservoir for clinically relevant MDR-GNB in need of increased surveillance, prominent differences in MDR-GNB composition suggest a need for complementary screening of both sites to capture the full spectrum of resistance within a healthcare system. Importantly, both bacterial strains and plasmids demonstrated longitudinal persistence and links to clinical cultures, highlighting the dual challenge of detecting indirect routes of MDR-GNB transmission via the environment and plasmids and supporting a role for comprehensive long-read sequencing to resolve transmission dynamics including those involving plasmid spread. Our work shows that environmental sampling and surveillance could play a key role in anticipatory screening for MDR-GNB, but that source-specific distribution of bacterial taxa and plasmids must be considered when interpreting the clinical relevance of environmental data. This reinforces the critical importance of developing multifaceted infection prevention strategies that include environmental surveillance, decontamination, and engineering controls to reduce environmental sources for MDR bacteria and reduce the clinical burden of antimicrobial resistance.

## Materials and Methods

### Environmental sample collection and screening

We performed environmental surveillance for MDR-GNB at a New York City hospital via sink drain surveillance swabs and wastewater testing (**Figure 1**). Sink drain swabs were collected from two inpatient hospital units (IHUA and IHUB) between March and September 2023 over five sampling timepoints (**Figure 1B**). For all timepoints, swabs were obtained from sink drains in all accessible patient rooms (including separate swabs for bathroom and handwashing sinks within the room if present) and the unit staff bathroom. Samples were taken at planned timepoints regardless of patient presence or daily or terminal cleaning status. During timepoints 1 and 2, we also swabbed high-touch surfaces within the hospital room (e.g. bed rails, side tables, IV pump interface) and unit nursing stations; however, this was not continued in timepoints 3 through 5 due to the low bacterial growth and MDR-GNB detection rates (10/73 (14%) high-touch swabs exhibited growth on non-selective media, and only 2/73 had MDR-GNB growth (3%)). Sink drain sampling was performed by inserting CultureSwabs sterile double-headed swabs (BD Diagnostic Systems) as far as possible into drain openings and vigorously swabbing the surface of the drainpipe inlet leading to the sink trap. One swab from each sample was used to inoculate a non-selective tryptic soy agar (TSA) plate, to establish that bacteria were successfully sampled from the site, and a CHROMagar ESBL plate for selective isolation of MDR-GNB. Inoculated plates were incubated overnight at 37°C. From each CHROMagar ESBL plate, up to two colonies per unique colony morphology (colony color, halo color, colony size) were subcultured onto CHROMagar KPC plates to identify whether each isolate exhibited resistance to extended-spectrum beta-lactams only (growth on CHROMagar ESBL and absence of growth on CHROMagar KPC) or also to carbapenems (growth on CHROMagar KPC). These phenotypes were used to categorize each isolate as third generation cephalosporin-resistant or carbapenem-resistant gram-negative bacteria (CephR-GNB, CR-GNB). Subculture plates were used to prepare purified bacterial isolates in TSB with 25% glycerol, which were stored at −80°C for downstream analyses.

Concomitant wastewater samples were collected as part of broader efforts related to viral and MDR bacterial hospital wastewater surveillance. As previously described,^26^ wastewater was collected from two house traps in the main hospital, referred to here as Hospital Quadrant A and B (HQA, HQB). HQA and HQB each collect wastewater from a vertical quadrant of the hospital building which include patient units IHUA and IHUB, respectively (i.e., wastewater from IHUA is captured by HQA and wastewater from IHUB is captured by HQB; **Figure 1A**). Composite (time-weighted, 24-hour) wastewater was collected weekly from each sampler for MDR-GNB surveillance. From each sample, 50 mL wastewater was briefly centrifuged (400 x g) to remove large debris. Microbes were then concentrated from wastewater using an InnovaPrep Concentrator Pipette using manufacturer recommended settings and eluted in PBS. Similar to the methods above for sink drain swabs, microbial concentrates were serially diluted from 10^0^ – 10^-5^, then plated on TSA plates to establish successful bacterial sampling and CHROMagar ESBL plates to isolate MDR-GNB. Up to two colonies per unique morphology were subcultured from CHROMagar ESBL onto CHROMagar KPC plates to determine each isolate’s final resistance phenotype (CephR-GNB vs CR-GNB) as defined above. Isolate stocks were prepared from subculture plates and stored at −80°C in TSB with 25% glycerol.

Study procedures were approved by the Columbia University Irving Medical Center Institutional Review Board (Protocol Nos. AAAT2294, AAAU1682).

### Whole genome sequencing

All environmental and wastewater MDR-GNB isolates underwent DNA extraction using the DNeasy PowerSoil Kit or DNeasy 96 PowerSoil Pro QIAcube HT Kit (Qiagen) with additional shearing and bead-based size selection to target DNA fragment lengths of ∼20 kb. We then performed nanopore long-read nanopore sequencing and assembly on all isolates in the collection to enable comprehensive genomic analysis and identification of plasmids and other MGEs harboring antibiotic resistance genes. Briefly, the Rapid Barcoding Kit (Oxford Nanopore Technologies) was used for library preparation and multiplexing (24-96 samples per run), with run times of 24-72 hours on GridION R9.4.1 or R10.4.1 flow cells (Oxford Nanopore Technologies).

Assemblies were generating using Flye v2.9,^36^ polished using Medaka v2.2,^37^ and annotated using Prokka v1.14^5^. Species identification was performed using both Mash v2.3^38^ and MLST v2.23^39^ on polished assemblies. ARG calling was performed using SRST2 for assemblies^40,41^ and StarAMR v0.10^42^ with the CARD v3.0.8^43^ database. Assembly quality metrics were assessed using CheckM2 v1.1^44^ and Bandage v0.9.^45^ Resulting assemblies were also visually curated using Bandage to ensure appropriate assembly size and contiguity. Criteria for inclusion in the final genomic dataset included CheckM2 completeness ≥ 90% and contamination ≤5%, Mash classification output with single species hit. We also excluded six isolates classified as gram-positive species, as these were likely artefactual given our selective culturing protocol (3 *Staphylococcus epidermidis*, 1 *Staphylococcus hominis*, 1 *Streptococcus oralis*, 1 *Streptococcus salivarius*). Final taxonomic classification (species and, if available, multilocus sequence type (ST)) were manually reviewed and curated as follows: MASH taxonomic classification was used as the default for species-level assignment after cross-referencing with the species-specific scheme selected by the MLST program. In almost all cases of disagreement, MASH was able to identify known subspecies within the species complex assigned by MLST (e.g., *Enterobacter hormaechei* assigned by MASH and *Enterobacter cloacae* complex scheme used by MLST); in these cases, the subspecies identification from MASH was used. We also searched available literature for up-to-date species or subspecies nomenclature for all STs assigned by MLST. For isolates assigned to species with available MLST schemes but unassigned to a known ST, allelic data from all genes in the relevant scheme were manually reviewed and grouped by similarity into novel ST assignments. Identical novel MLST profiles and any single locus variants were grouped together as a single novel ST.

As described above, up to two colonies per morphology were isolated from each original inoculation plate to reasonably capture species and strain diversity. We implemented a stepwise strategy to prevent bias in our dataset due to this isolation protocol when calculating species, strain, gene, or plasmid-level diversity based on collection location. For each sink drain or wastewater quadrant on a given collection date, isolates with identical species, sequence type, and ARG profiles were grouped together; the following stepwise criteria were used to select a single representative isolate and exclude others from further analysis: highest completeness, then lowest level of contamination, then highest coverage of longest contig, then longest contig length, then lowest total contig number.

### Spatiotemporal analysis

We analyzed longitudinal trends in MDR-GNB across environmental sources (sink drains in inpatient hospital units versus hospital wastewater) and specific collection sites (IHUA, IHUB, HQA, HQB) using several methods. First, we quantified the prevalence of various MDR species at each environmental collection site and across sampling timepoints. Separately for CephR- and CR-GNB, we calculated the proportion of sampled sinks or wastewater samples from which each species was isolated at a given site and timepoint. Next, we performed a more granular analysis to establish the relatedness of sampling sites based on the prevalence of specific CephR or CR STs. Pearson distance was used to cluster heatmap rows (unique STs) and columns (sampling site and timepoint) to test for clustering by sampling source (sink drains versus wastewater), site, and/or timepoint. Lastly, for each timepoint, we summarized the proportion of unique MDR-GNB collected at each sampling site that were assigned to a given species and also identified the number of shared versus distinct STs between sink drains and wastewater, and between sites within each environmental source (i.e., IHUA vs IHUB, HQA vs HQB).

### ARG and plasmid analysis

ARG alleles present in each isolate were summarized using SRST2 outputs. For environmental and wastewater MDR-GNB isolates harboring *bla*_CTX-M_, *bla*_KPC_, and *bla*_NDM_, we also extracted resistance gene-encoding plasmid sequences from whole genome assemblies using StarAMR and Seqkit to assess plasmid diversity and sharing across isolate sources.

Our plasmid genotyping and clustering approach has been described previously.^14^ Briefly, plasmid genotyping (e.g. replicon and relaxase typing) and mobility prediction (conjugative, mobilizable, or non-mobilizable) were performed using MOB-typer.^46^ Contigs greater than 400 kb in length and/or lacking any plasmid genotypic markers were excluded from plasmid clustering analysis. We then used MOB-cluster to create a closed plasmid reference database and identify closely related plasmids within ARG-specific datasets based on Mash distance cutoffs (0.06 for primary clusters and 0.025 for secondary clusters).

To visualize the movement of ARGs and ARG-harboring plasmids over sampling timepoints and within the hospital, we generated a representation of inpatient hospital unit layouts and wastewater quadrants. For each sampling timepoint, antibiotic resistance alleles and plasmid secondary clusters were plotted using *ggimage* in R based on the original sink drain or wastewater location where each isolate was collected.

### Clinical MDR-GNB collection

Previously assembled CRE and CephRE genomes of clinical isolates from the same hospital center were compared to environmental isolates in this study. These included 164 CTX-M-producing CephRE isolates collected between June 2015-January 2019 (BioProject PRJNA1191487), 435 KPC-producing CRE isolates collected between 2009-2018 (BioProjects PRJNA1088550 and PRJNA759273), and 3 NDM-producing CRE collected from March-April 2020 (BioProject PRJNA645930).^14,20^

### Links across environmental and clinical MDR-GNB collections

For each genus with at least two isolates in our study, including both environmental and clinical collections (13 of 18 genera total), we generated similarity matrices between all isolate pairs based on pairwise average nucleotide identity (ANI). Chromosomal contigs were extracted from each assembly using Platon v1.7^47^ and used as the input for FastANI v1.34.^48^ FastANI matrices were then used to create network plots using *ggraph*^49^ in R. Network clusters were defined using a conservative threshold of ANI>99.5%, based on inspection of pairwise distance density histograms and dendrograms for each genus generated using *bactaxR*.^50^

We also assessed plasmid relatedness between *bla*_CTX-M_, *bla*_KPC_, and *bla*_NDM_*-*harboring plasmids from environmental isolates in this study and the prior genomic studies of clinical CRE and ESBLE isolates described above. MOB-cluster was used to generate new closed plasmid reference databases containing both environmental and clinical resistance plasmids, and to identify plasmid clusters based on Mash distance cutoffs.

To identify links between MDR-GNB across collections, shared genomes were defined as genomes with ANI>99.5%, and shared plasmids were defined as belonging to the same secondary MOB-cluster (which applies Mash with complete-linkage clustering at a Mash distance cutoff of 0.025). All unique isolate pairs in this study which had shared genomes, plasmids, or both, as defined above, were identified. We analyzed both the complete dataset of all isolate pairs as well as the subset of pairs where isolates were collected from two different collections (e.g., HQA vs HQB) to assess chromosome or plasmid sharing within and across collection sites. The R library *circlize*^51^ was used to generate a circular plot visualizing the number of genomic, plasmid, or combined links between each set of sources.

### Statistical Analyses

Chi-squared tests for independence (implemented using chisq.test() in R) were used to test for associations between culture-based detection of CephR- and CR-GNB from the same sink drain (**Figure S3**). Wilcoxon tests were used to compare pairwise ANI values across isolate pairs based on collection location (**Figure 3**, **Figure S6**). Lastly, two-proportion Z-tests (implemented using prop.test() in R) and Chi-squared goodness-of-fit tests (implemented using chisq.test() in R) were also used to test whether links between environmental and clinical datasets were driven primarily by sharing of bacterial strains versus resistance plasmids, and by sharing between clinical isolate and sink drains versus wastewater.

## Data availability

All isolate genomes were deposited to the NCBI Genome repository (https://www.ncbi.nlm.nih.gov/genome/) under BioProject numbers PRJNA1503005 (sink drain isolates) and PRJNA1502998 (wastewater isolates). Deidentified collection data and genotypic metadata for each isolate is publicly available in the BioSample records listed under these two BioProjects. As described in the main text, previously published genomes of clinical isolates used for comparison in this study are also available at BioProjects PRJNA1191487, PRJNA1088550, PRJNA759273, and PRJNA645930.

## Acknowledgments

M.K.A and A.G.-S. conceived of and designed the study with input from A.-C.U. M.K.A., T.W.H., and A.G.-S. performed hospital unit sampling, and M.K.A. carried out wastewater sampling. M.K.A. and T.W.H. performed bacterial culture, DNA extraction, and whole genome sequencing with assistance from A.G.S. The bioinformatics pipeline was developed and implemented by M.K.A., who analyzed the data together with A.G.-S., and K.T. M.K.A and A.G.-S. wrote the manuscript, with critical support from all authors. All authors read and approved the submitted version.

## References

1. Centers for Disease Control and Prevention (U.S.). Antibiotic Resistance Threats in the United States, 2019. https://stacks.cdc.gov/view/cdc/82532 (2019).

2. Jernigan, J. A. et al. Multidrug-Resistant Bacterial Infections in U.S. Hospitalized Patients, 2012-2017. N. Engl. J. Med. 382, 1309–1319 (2020).

3. Blanco, N., O’Hara, L. M. & Harris, A. D. Transmission pathways of multidrug-resistant organisms in the hospital setting: a scoping review. Infect. Control Hosp. Epidemiol. 40, 447–456 (2019).

4. Chia, P. Y. et al. The role of hospital environment in transmissions of multidrug-resistant gram-negative organisms. Antimicrob. Resist. Infect. Control 9, 29 (2020).

5. Marimuthu, K. et al. Whole genome sequencing reveals hidden transmission of carbapenemase-producing Enterobacterales. Nat. Commun. 13, 3052 (2022).

6. Sauerborn, E. et al. Resolving plasmid-encoded carbapenem resistance dynamics and reservoirs in a hospital setting through nanopore sequencing. *Microb*. Genomics 12, (2026).

7. Kanamori, H., Weber, D. J. & Rutala, W. A. Healthcare Outbreaks Associated With a Water Reservoir and Infection Prevention Strategies. Clin. Infect. Dis. Off. Publ. Infect. Dis. Soc. Am. 62, 1423–1435 (2016).

8. Kizny Gordon, A. E., et al. The Hospital Water Environment as a Reservoir for Carbapenem-Resistant Organisms Causing Hospital-Acquired Infections-A Systematic Review of the Literature. Clin. Infect. Dis. Off. Publ. Infect. Dis. Soc. Am. 64, 1435–1444 (2017).

9. Volling, C. et al. Are Sink Drainage Systems a Reservoir for Hospital-Acquired Gammaproteobacteria Colonization and Infection? A Systematic Review. Open Forum Infect. Dis. 8, ofaa590 (2021).

10. Regev-Yochay, G. et al. Sink-traps are a major source for carbapenemase-producing *Enterobacteriaceae* transmission. Infect. Control Hosp. Epidemiol. 45, 284–291 (2024).

11. Anantharajah, A. et al. Long-term intensive care unit outbreak of carbapenemase-producing organisms associated with contaminated sink drains. J. Hosp. Infect. 143, 38–47 (2024).

12. Kotay, S., Chai, W., Guilford, W., Barry, K. & Mathers, A. J. Spread from the Sink to the Patient: *In Situ* Study Using Green Fluorescent Protein (GFP)-Expressing Escherichia coli To Model Bacterial Dispersion from Hand-Washing Sink-Trap Reservoirs. Appl. Environ. Microbiol. 83, e03327–16 (2017).

13. Chng, K. R. et al. Cartography of opportunistic pathogens and antibiotic resistance genes in a tertiary hospital environment. Nat. Med. 26, 941–951 (2020).

14. Gomez-Simmonds, A. et al. Genomic epidemiology of carbapenem-resistant Enterobacterales at a New York City hospital over a ten-year period reveals complex plasmid-clone dynamics and evidence for frequent horizontal transfer of blaKPC. Genome Res. gr.279355.124 (2024) doi:10.1101/gr.279355.124.

15. Koh, V. et al. Plasmid dynamics driving carbapenemase gene dissemination in healthcare environments: a nationwide analysis of closed Enterobacterales genomes. Nat. Commun. 16, 9522 (2025).

16. Roberts, L. W. et al. Long-read sequencing reveals genomic diversity and associated plasmid movement of carbapenemase-producing bacteria in a UK hospital over 6 years. *Microb*. Genomics 9, (2023).

17. Macesic, N. et al. Genomic investigation of multispecies and multivariant *bla*_NDM_ outbreak reveals key role of horizontal plasmid transmission. Infect. Control Hosp. Epidemiol. 45, 709–716 (2024).

18. McCallum, G. E. & Hall, J. P. J. The hospital sink drain microbiome as a melting pot for AMR transmission to nosocomial pathogens. Npj Antimicrob. Resist. 3, 68 (2025).

19. Gomez-Simmonds, A. et al. Population structure of blaKPC-harbouring IncN plasmids at a New York City medical centre and evidence for multi-species horizontal transmission. J. Antimicrob. Chemother. 77, 1873–1882 (2022).

20. Gomez-Simmonds, A. et al. Carbapenemase-producing Enterobacterales causing secondary infections during the COVID-19 crisis at a New York City hospital. J. Antimicrob. Chemother. 76, 380–384 (2021).

21. Gomez-Simmonds, A. et al. Genomic and Geographic Context for the Evolution of High-Risk Carbapenem-Resistant Enterobacter cloacae Complex Clones ST171 and ST78. mBio 9, e00542–18 (2018).

22. Weingarten, R. A. et al. Genomic Analysis of Hospital Plumbing Reveals Diverse Reservoir of Bacterial Plasmids Conferring Carbapenem Resistance. mBio 9, e02011–17 (2018).

23. Carlsen, L. et al. Long time persistence and evolution of carbapenemase-producing Enterobacterales in the wastewater of a tertiary care hospital in Germany. J. Infect. Public Health 16, 1142–1148 (2023).

24. Diorio-Toth, L. et al. Intensive care unit sinks are persistently colonized with multidrug resistant bacteria and mobilizable, resistance-conferring plasmids. mSystems e00206–23 (2023) doi:10.1128/msystems.00206-23.

25. Greendyke, W., Mantell, E., Moore, E., Cave, E. & Kratz, M. Creation and Distribution of 2021–2023 New York City Regional Antibiograms. Antimicrob. Steward. Healthc. Epidemiol. 5, s129–s129 (2025).

26. Annavajhala, M. K. et al. Hospital wastewater surveillance for SARS-CoV-2 identifies intra-hospital dynamics of viral transmission and evolution. Appl. Environ. Microbiol. 91, e00501–25 (2025).

27. Tsang, K. K. et al. Diversity, functional classification and genotyping of SHV β-lactamases in Klebsiella pneumoniae. *Microb*. Genomics 10, (2024).

28. Ledwoch, K., Robertson, A., Lauran, J., Norville, P. & Maillard, J.-Y. It’s a trap! The development of a versatile drain biofilm model and its susceptibility to disinfection. J. Hosp. Infect. 106, 757–764 (2020).

29. Sukhum, K. V. et al. Antibiotic-resistant organisms establish reservoirs in new hospital built environments and are related to patient blood infection isolates. Commun. Med. 2, 62 (2022).

30. Shams, A. M. et al. Assessment of the Overall and Multidrug-Resistant Organism Bioburden on Environmental Surfaces in Healthcare Facilities. Infect. Control Hosp. Epidemiol. 37, 1426–1432 (2016).

31. Qiao, F. et al. Handwashing Sink Contamination and Carbapenem-resistant *Klebsiella* Infection in the Intensive Care Unit: A Prospective Multicenter Study. Clin. Infect. Dis. 71, S379–S385 (2020).

32. La Rosa, M. C. et al. The Impact of Wastewater on Antimicrobial Resistance: A Scoping Review of Transmission Pathways and Contributing Factors. Antibiotics 14, 131 (2025).

33. D’Souza, A. W. et al. Spatiotemporal dynamics of multidrug resistant bacteria on intensive care unit surfaces. Nat. Commun. 10, 4569 (2019).

34. Bourdin, T. et al. Disinfection of sink drains to reduce a source of three opportunistic pathogens, during Serratia marcescens clusters in a neonatal intensive care unit. PLOS ONE 19, e0304378 (2024).

35. Gomez-Simmonds, A. et al. Population structure of *bla* KPC-harbouring IncN plasmids at a New York City medical centre and evidence for multi-species horizontal transmission. J. Antimicrob. Chemother. 77, 1873–1882 (2022).

36. Kolmogorov, M., Yuan, J., Lin, Y. & Pevzner, P. A. Assembly of long, error-prone reads using repeat graphs. Nat. Biotechnol. 37, 540–546 (2019).

37. Medaka. Oxford Nanopore Technologies.

38. Ondov, B. D. et al. Mash: fast genome and metagenome distance estimation using MinHash. Genome Biol. 17, 132 (2016).

39. Seemann, T. mlst.

40. Wick, R. SRST2 table from assemblies.

41. Inouye, M. et al. SRST2: Rapid genomic surveillance for public health and hospital microbiology labs. Genome Med. 6, 90 (2014).

42. Bharat, A. et al. Correlation between Phenotypic and In Silico Detection of Antimicrobial Resistance in Salmonella enterica in Canada Using Staramr. Microorganisms 10, (2022).

43. Alcock, B. P. et al. CARD 2023: expanded curation, support for machine learning, and resistome prediction at the Comprehensive Antibiotic Resistance Database. Nucleic Acids Res. 51, D690–D699 (2023).

44. Chklovski, A., Parks, D. H., Woodcroft, B. J. & Tyson, G. W. CheckM2: a rapid, scalable and accurate tool for assessing microbial genome quality using machine learning. Nat. Methods 20, 1203–1212 (2023).

45. Wick, R. R., Schultz, M. B., Zobel, J. & Holt, K. E. Bandage: interactive visualization of de novo genome assemblies. Bioinformatics 31, 3350–3352 (2015).

46. Robertson, J. & Nash, J. H. E. MOB-suite: software tools for clustering, reconstruction and typing of plasmids from draft assemblies. *Microb*. Genomics 4, (2018).

47. Schwengers, O. et al. Platon: identification and characterization of bacterial plasmid contigs in short-read draft assemblies exploiting protein sequence-based replicon distribution scores. *Microb*. Genomics 6, (2020).

48. Jain, C., Rodriguez-R, L. M., Phillippy, A. M., Konstantinidis, K. T. & Aluru, S. High throughput ANI analysis of 90K prokaryotic genomes reveals clear species boundaries. Nat. Commun. 9, 5114 (2018).

49. Pedersen, T. ggraph: An Implementation of Grammar of Graphics for Graphs and Networks.

50. Carroll, L. M., Wiedmann, M. & Kovac, J. Proposal of a Taxonomic Nomenclature for the Bacillus cereus Group Which Reconciles Genomic Definitions of Bacterial Species with Clinical and Industrial Phenotypes. mBio 11, 10.1128/mbio.00034-20 (2020).

51. Gu, Z., Gu, L., Eils, R., Schlesner, M. & Brors, B. *circlize* implements and enhances circular visualization in R. Bioinformatics 30, 2811–2812 (2014).

