## Supplemental Figure 1 for "Interconnected reservoirs of multidrug-resistant bacteria and plasmids within hospital plumbing infrastructure"

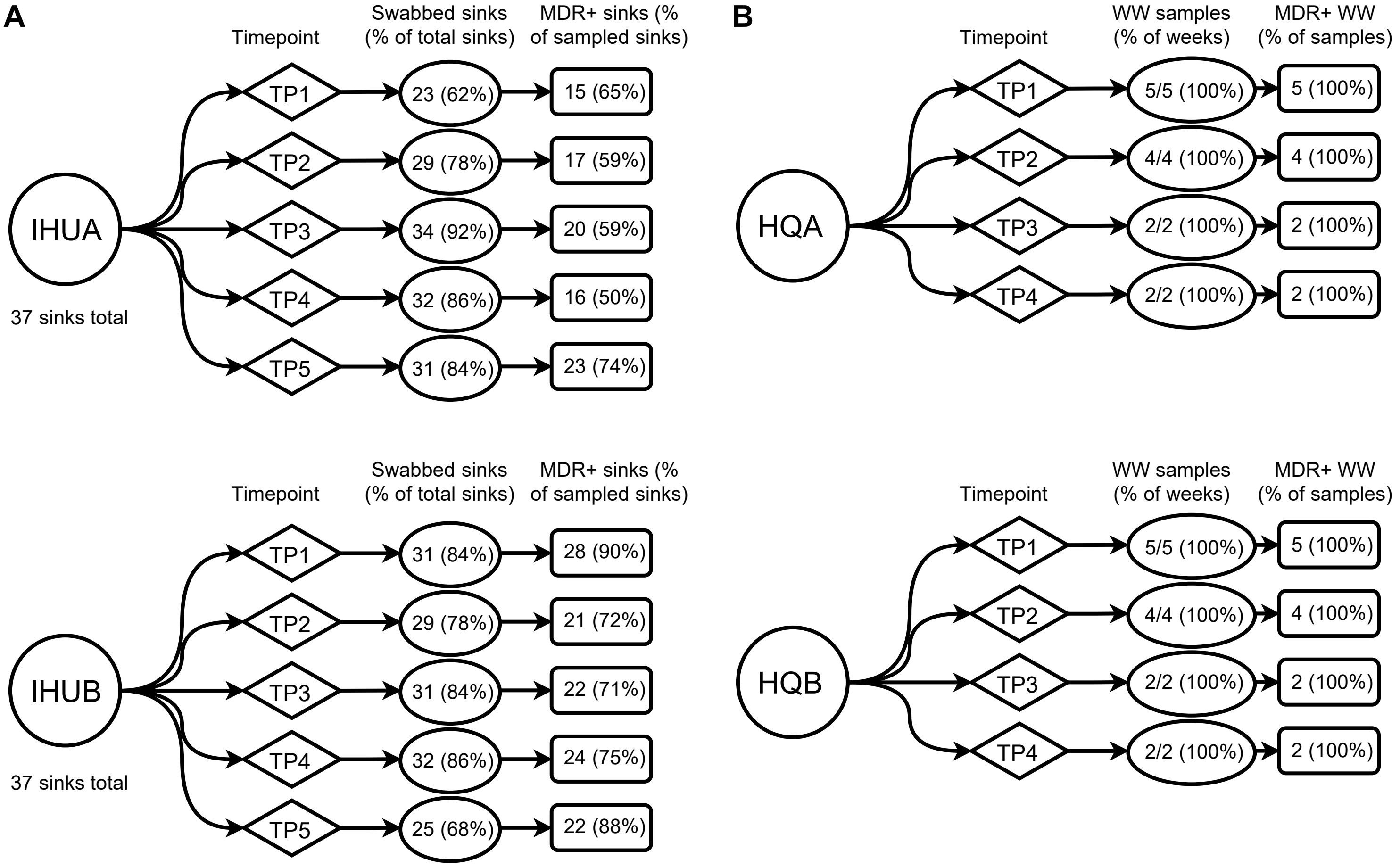


**Figure S1. High prevalence of multidrug-resistant bacteria in the hospital plumbing environment. (A)** Sinks swabs were collected from hospital rooms and bathrooms in two inpatient hospital units, IHUA and IHUB, over five different timepoints between March 2023 and September 2023 (timepoints TP1-TP5, shown as diamonds). A total of 37 different sinks were available for swabbing in each unit; the fraction of sinks sampled at each timepoint is shown and ranged from 62-92%. Multidrug-resistant Gram-negative bacteria (MDR-GNB) were highly prevalent, with 50-74% of sampled sinks in IHUA and 71-90% of sampled sinks in IHUB identified as MDR-GNB-positive (MDR+) based on selective culture. **(B)** Hospital wastewater from hospital quadrants A and B, overlapping with IHUA and IHUB, respectively (see Figure S1), was collected during concomitant timepoints for broader pathogen surveillance efforts. Wastewater from both quadrants was universally positive for MDR-GNB, further highlighting their prevalence in hospital plumbing infrastructure.
