## Supplemental Figure 2 for "Interconnected reservoirs of multidrug-resistant bacteria and plasmids within hospital plumbing infrastructure"

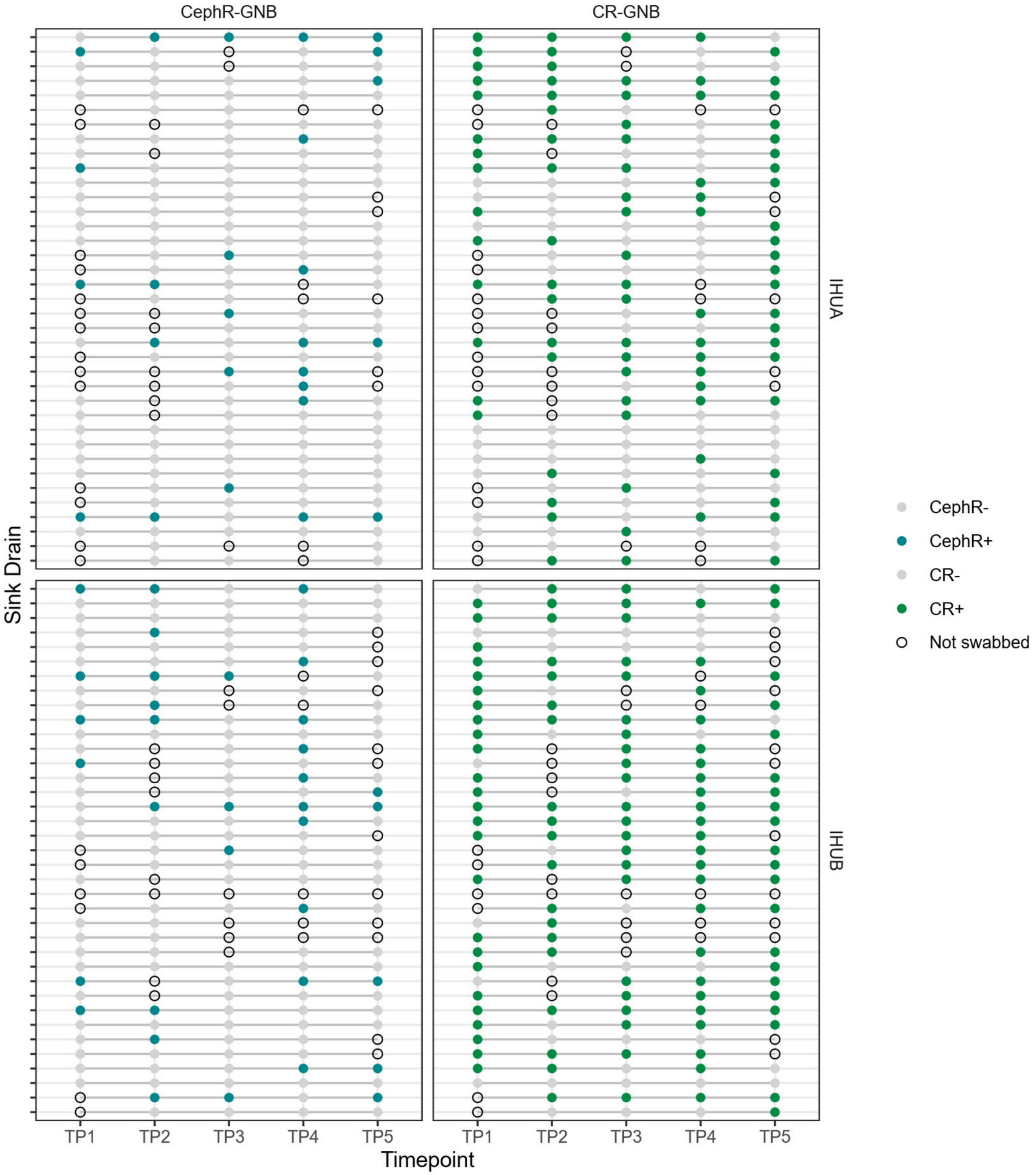


**Figure S2. MDR-GNB growth from sink drain swabs.** Growth of third generation cephalosporin- or carbapenem-resistant gram-negative bacteria (CephR-GNB, CR-GNB) was assessed via selective culture for each sink drain (rows) in inpatient hospital units IHUA and IHUB over the five sampling timepoints TP1-5. Each grey or colored dot represents a collected sink drain swab; teal or green dots indicate swabs positive for CephR- or CR-GNB, respectively. Sink drains not swabbed at a specific timepoint are shown as open circles. CephR-GNB grew during at least one timepoint in 15 (41%) sinks in IHUA and 19 (51%) of sinks in IHUB; CR-GNB grew during at least one timepoint in 34 (92)% of IHUA sinks and 34 (92%) of IHUB sinks. CephR-GNB were persistent in 5 (14%) of the sink drains in IHUA and 8 (22%) of the sink drains in IHUB, defined as growth of a given phenotype over two or more consecutive timepoints. CR-GNB were persistent in 20 (54%) IHUA sinks and 27 (73%) IHUB sinks.
