## Supplemental Figure 3 for "Interconnected reservoirs of multidrug-resistant bacteria and plasmids within hospital plumbing infrastructure"

### All Sinks (IHUA + IHUB)

| | CR-GNB+ | CR-GNB- | |
|---|---|---|---|
| CephR-GNB+ | 48 | 12 | N=60 |
| CephR-GNB- | 148 | 89 | N=237 |
| | N=196 | N=101 | |

$X^2$ (df=1, N=297 total swabs): 5.8, p=0.02

### IHUA

| | CR-GNB+ | CR-GNB- | |
|---|---|---|---|
| CephR-GNB+ | 21 | 5 | N=26 |
| CephR-GNB- | 65 | 58 | N=123 |
| | N=86 | N=63 | |

$X^2$ (df=1, N=149 total swabs): 5.8, p=0.02

### IHUB

| | CR-GNB+ | CR-GNB- | |
|---|---|---|---|
| CephR-GNB+ | 27 | 7 | N=34 |
| CephR-GNB- | 83 | 31 | N=114 |
| | N=110 | N=38 | |

$X^2$ (df=1, N=148 total swabs): 0.32, p=0.58

**Figure S3. Relationship between CephR- and CR-GNB growth in hospital sink drains.** Chi-squared tests were performed to assess whether CephR-GNB and CR-GNB growth were significantly associated in all sink drains and separately for sinks in IHUA and IHUB. In the complete dataset, isolation of the two phenotypes were significantly associated for a given sink drain ($X^2$ (df=1, N=297 total swabs): 5.8, p=0.02). Further investigation for each hospital unit revealed that this effect was driven by sink drains in IHUA, where growth of CephR- and CR-GNB was strongly associated ($X^2$ (df=1, N=149 total swabs): 5.8, p=0.02). In contrast, this association was absent in IHUB sink drains $X^2$ (df=1, N=148 total swabs): 0.32, p=0.58).
