## Supplemental Figure 4 for "Interconnected reservoirs of multidrug-resistant bacteria and plasmids within hospital plumbing infrastructure"

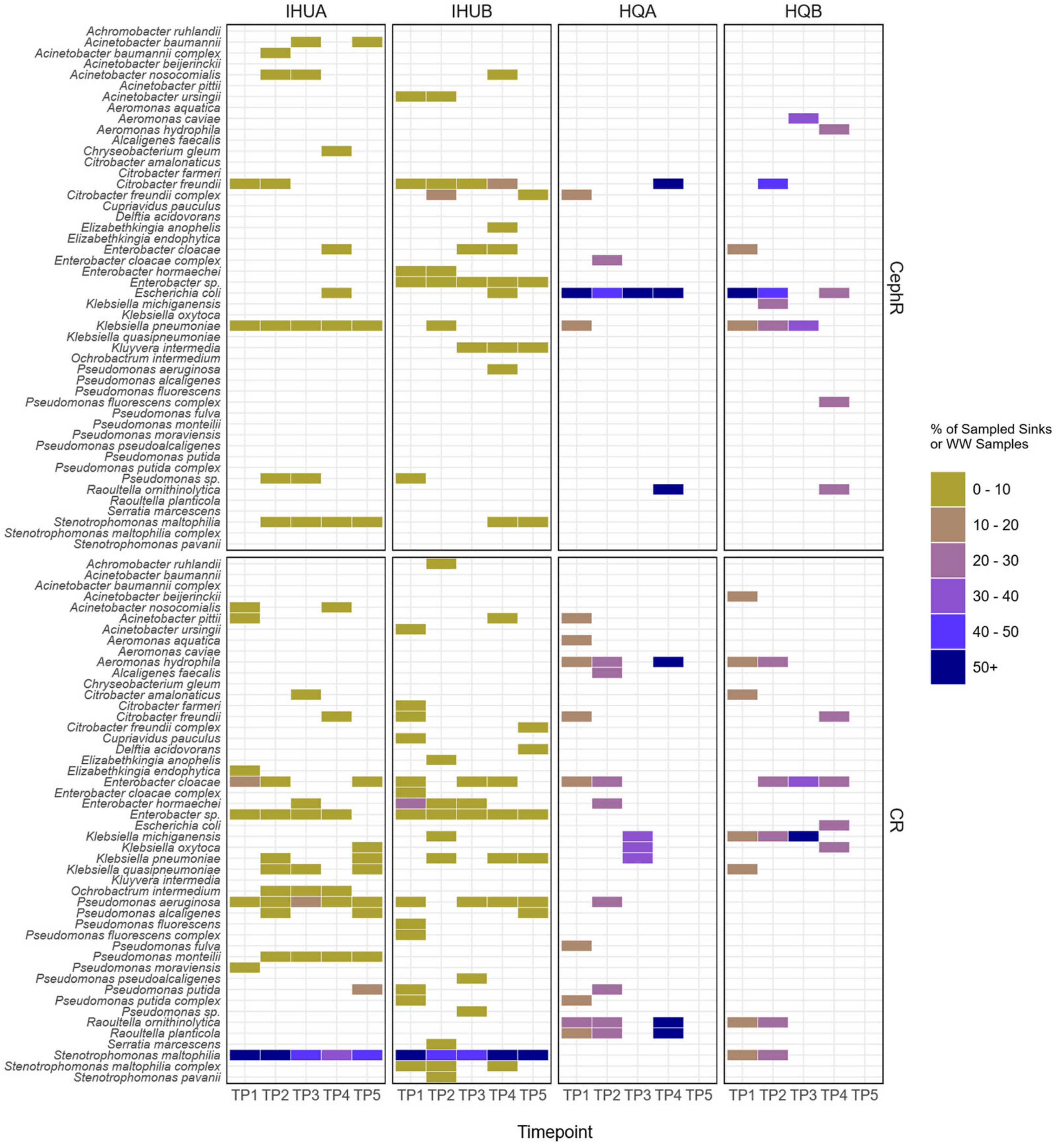


**Figure S4. Species prevalence across sampling timepoints.** For IHUA and IHUB, color intensity indicates the percent of sampled sinks at a given timepoint which grew each species. For HQA and HQB, color intensity indicates the percent of collected wastewater samples for a given timepoint which grew each species. For each species, data was aggregated separately for isolates with CephR- versus CR-GNB phenotypes, as shown in the top and bottom panels. *E. coli* dominated wastewater CephR-GNB at both sites, though *Klebsiella pneumoniae* also persisted in HQB over several consecutive timepoints. CR-GNB taxa were more distinct across the two wastewater quadrants; several *Klebsiella* and *Raoultella* species contributed to the majority of HQA CR-GNB burden over time, while *Enterobacter cloacae* and *Klebsiella michiganensis* more consistently drove CR-GNB detection in HQB. In sink drains, across all timepoints, diverse species of CephR-GNB were isolated, with Klebsiella pneumoniae isolated at all timepoints in IHUA while Enterobacter sp. was isolated at all timepoints in IHUB. CR-GNB in sink drains from both inpatient units was dominated by *Stenotrophomonas maltophilia,* with over 30% of sampled sink drains at all timepoints growing CR-*S. maltophilia*.
