## Supplemental Figure 5 for "Interconnected reservoirs of multidrug-resistant bacteria and plasmids within hospital plumbing infrastructure"

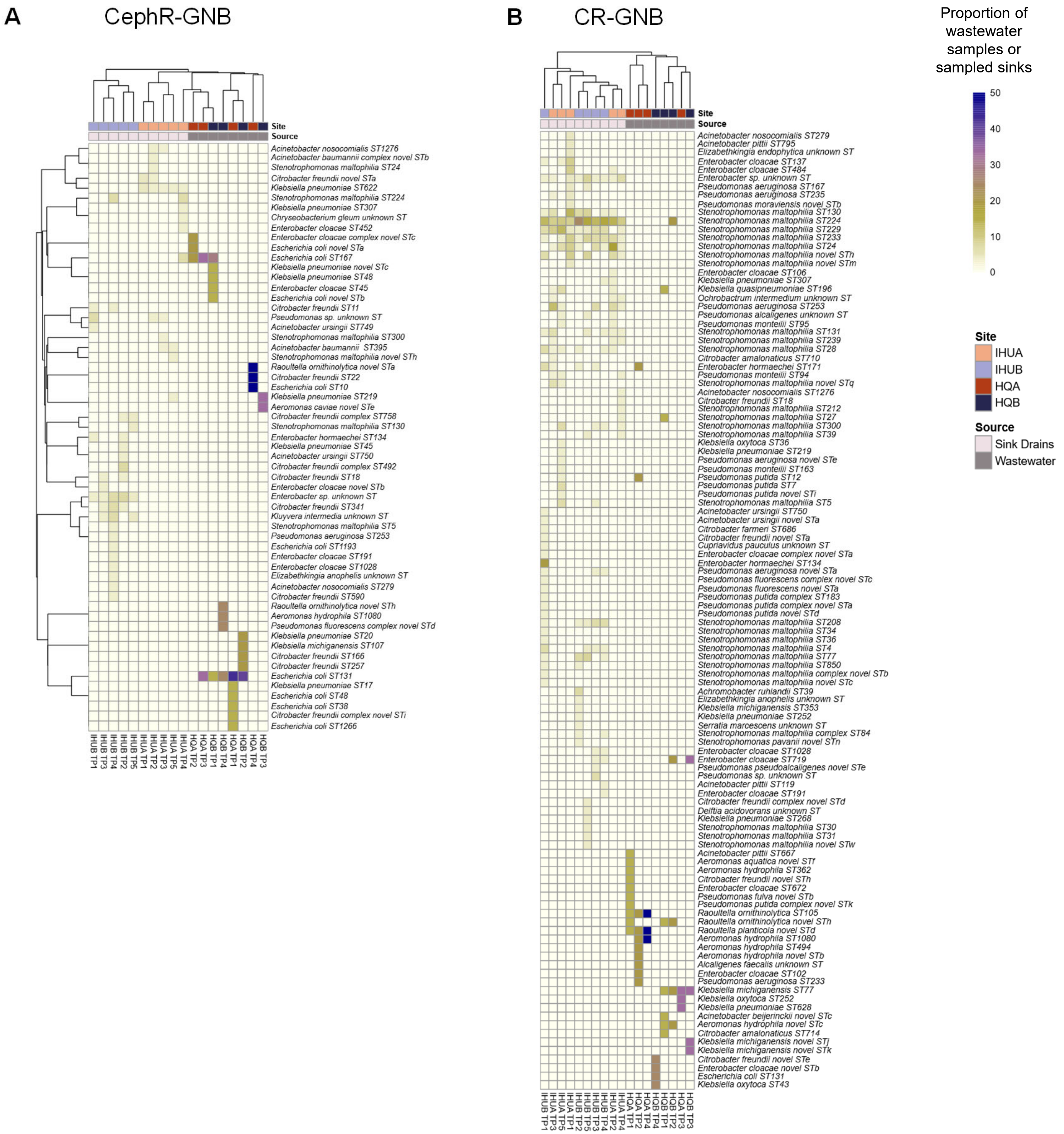


**Figure S5. Clustering of environmental sites and sampling timepoints based on bacterial strain prevalence.** For each sampling timepoint, we calculated the proportion of wastewater samples or sampled sinks that were positive for each sequence type (ST) identified in environmental CephR- (panel **A**) or CR-GNB (panel **B**). These proportions were then used to generate heatmaps where sampling sites and timepoints (columns) and STs (rows) were clustered based on Pearson distances. For both CephR- and CR-GNB, hospital inpatient units IHUA and IHUB and wastewater quadrants HQA and HQB primarily clustered by environmental source (sink drains versus wastewater). Across all timepoints, collection sites belonging to the same environmental source (sink drains or wastewater) demonstrated similar CephR- and CR-GNB ST profiles. IHUA and IHUB clustered together and were distinct from HQA and HQB for both CephR- and CR-GNB. In sink drains, CephR- also clustered by collection site across timepoints (IHUA timepoints were distinct from IHUB timepoints). Overall, diverse CephR- and CR-GNB STs were recovered from sink drains across timepoints, while wastewater sites were characterized by dominance of a small number of CephR- or CR-GNB STs.
