## Supplemental Figure 6 for "Interconnected reservoirs of multidrug-resistant bacteria and plasmids within hospital plumbing infrastructure"

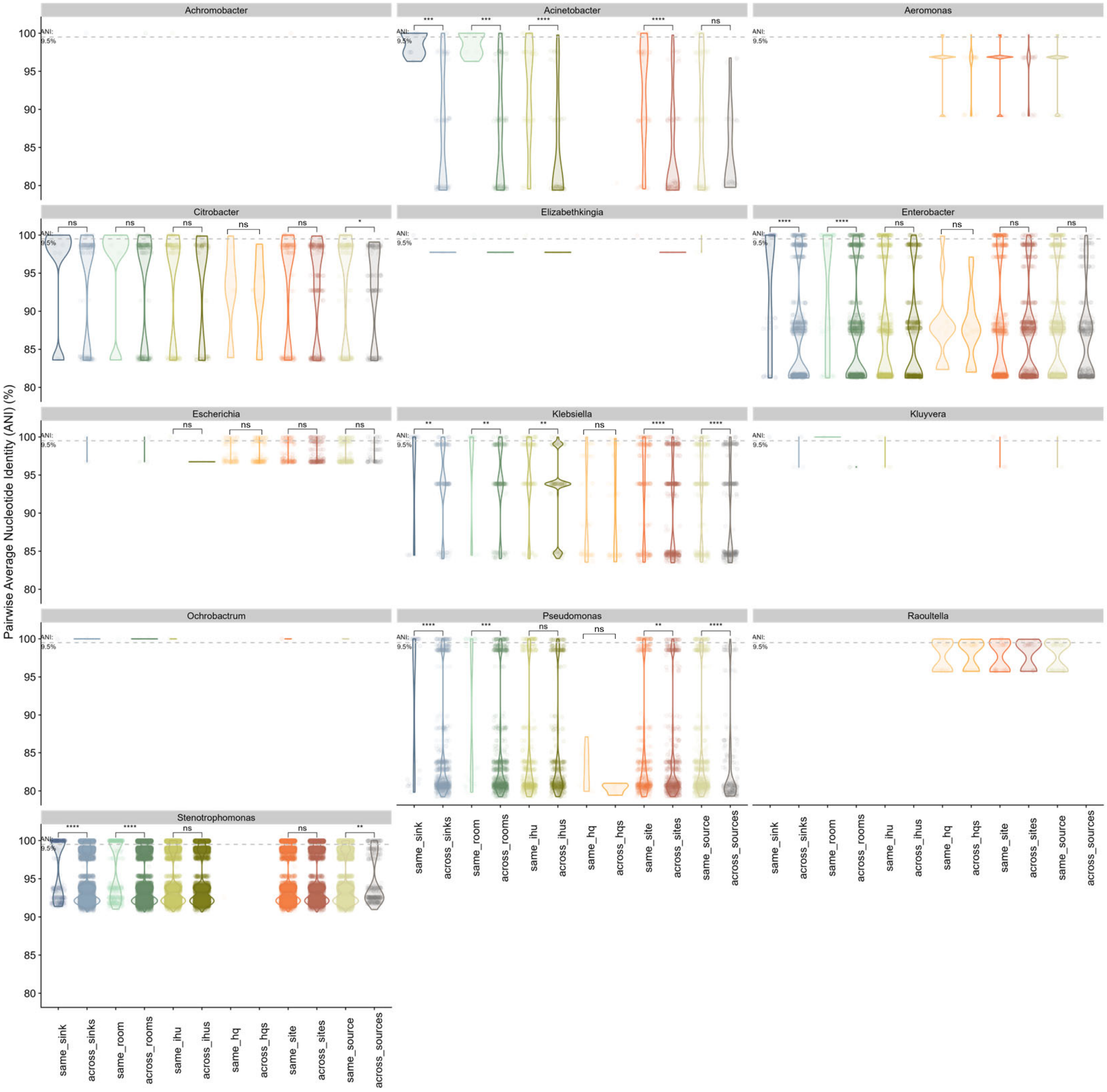


**Figure S6. Genomic relatedness of environmental MDR-GNB isolates by genus.** Pairwise average nucleotide identity (ANI) was calculated for all sink drain and wastewater isolates belonging to the same genus. Data was then categorized based on isolate collection source and compared, for each genus, across categories using Wilcox tests. Isolates from the same sink (at any collection timepoint) were compared to sink drain isolate pairs from different sinks (blue); to account for patient rooms with multiple sink drains, isolates collected from sink drains in the same patient room were also compared to those from different patient rooms (teal). Isolates from sink drains within the same inpatient hospital unit (IHU) were compared to those from different IHUs (i.e., IHUA vs IHUB; olive green), as were isolates the same hospital quadrant (HQ) wastewater compared to those from different HQs (i.e., HQA vs HQB; peach). MDR-GNB pairs from the same collection site (i.e., within IHUA, IHUB, HQA, or HQB; orange) and from the same environmental source (i.e., within sink drains or wastewater; beige) were also compared to those from different locations (maroon) or environments (grey). For each plot; a grey dashed horizontal line indicates the ANI 99.5% threshold. (* = p<0.05; ** = p<0.01; *** = p<0.001; **** = p<0.0001 based on Wilcoxon tests)
