## Supplemental Figure 7 for "Interconnected reservoirs of multidrug-resistant bacteria and plasmids within hospital plumbing infrastructure"

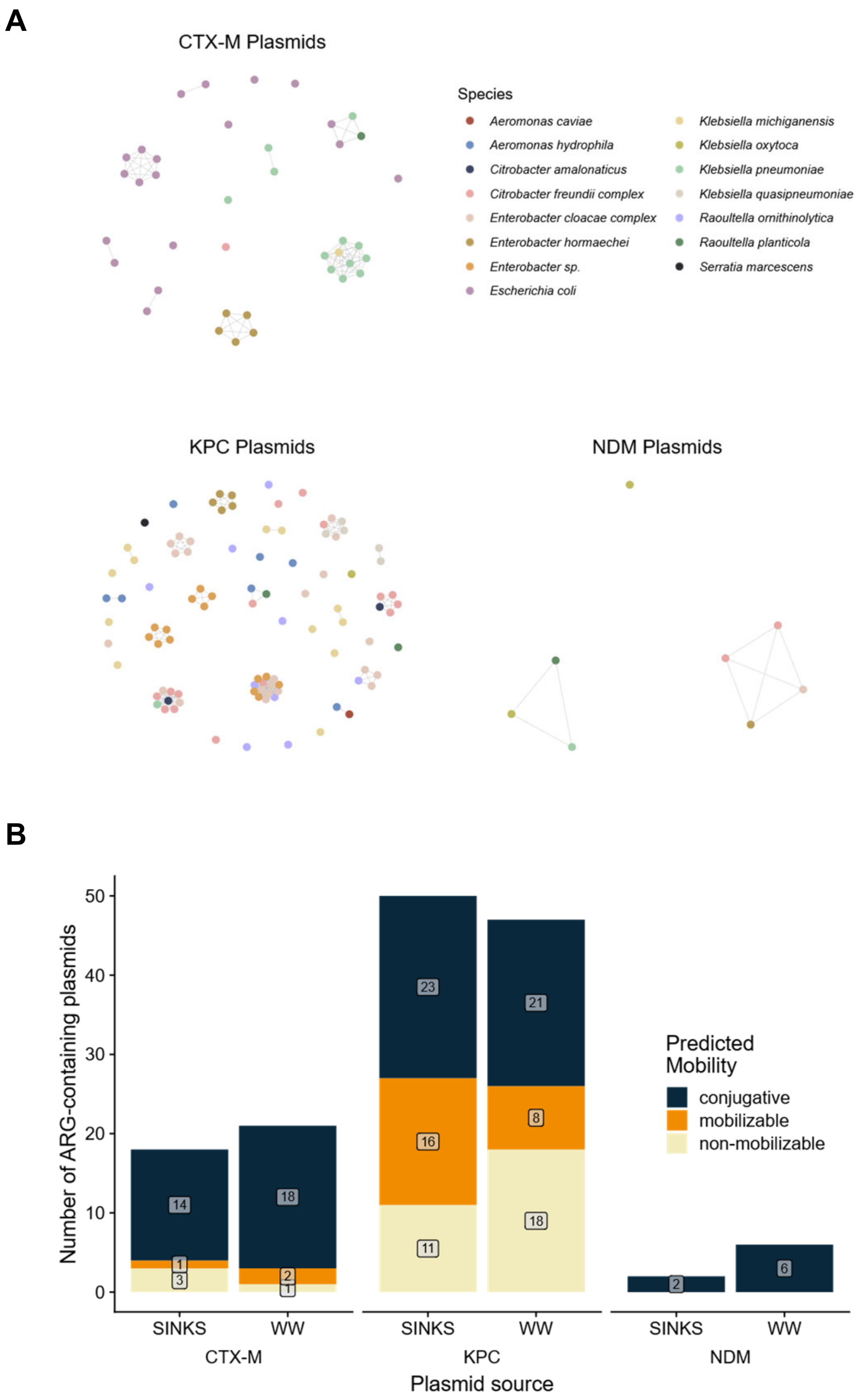


**Figure S7. Antibiotic resistance gene plasmids harbored by environmental MDR-GNB.** (**A**) Network plots based on plasmid sequence similarity as calculated using Mash distances by MOB cluster. Each colored point represents a plasmid harboring *bla*$_{\text{CTX-M}}$, *bla*$_{\text{KPC}}$, or *bla*$_{\text{NDM}}$. Colors indicate the species of the bacterial host from which the plasmid was isolated. Grey lines connecting dots define plasmid secondary clusters based on a Mash distance threshold of 0.025. *bla*$_{\text{CTX-M}}$-harboring plasmids formed 15 secondary clusters, including 2 containing plasmids found in at least two different species: one cluster identified in *K. pneumoniae* and *K. michiganensis* isolates, and another harbored by *K. pneumoniae*, *R. planticola*, and *E. coli*. *bla*$_{\text{KPC}}$ was harbored by plasmids from 42 secondary clusters, 7 of which comprised plasmids found in multiple species within the Enterobacterales family. 2 of the 3 plasmid secondary clusters harboring *bla*$_{\text{NDM}}$ contained plasmids found in three distinct Enterobacterales species. (**B**) Predicted mobility of ARG-harboring plasmids in environmental isolates based on MOB typer. Almost all *bla*$_{\text{CTX-M}}$ plasmids were predicted to be conjugative or mobilizable (35/39 *bla*$_{\text{CTX-M}}$ plasmids, including 15/18 from sink drains and 20/21 from wastewater), as were the majority of *bla*$_{\text{KPC}}$ plasmids (68/97 plasmids, including 39/50 from sink drains and 29/47 from wastewater). All 8 *bla*$_{\text{NDM}}$ plasmids, 2 from sink drains and 6 from wastewater, were predicted to be conjugative.
