## Supplemental Figure 8 for "Interconnected reservoirs of multidrug-resistant bacteria and plasmids within hospital plumbing infrastructure"

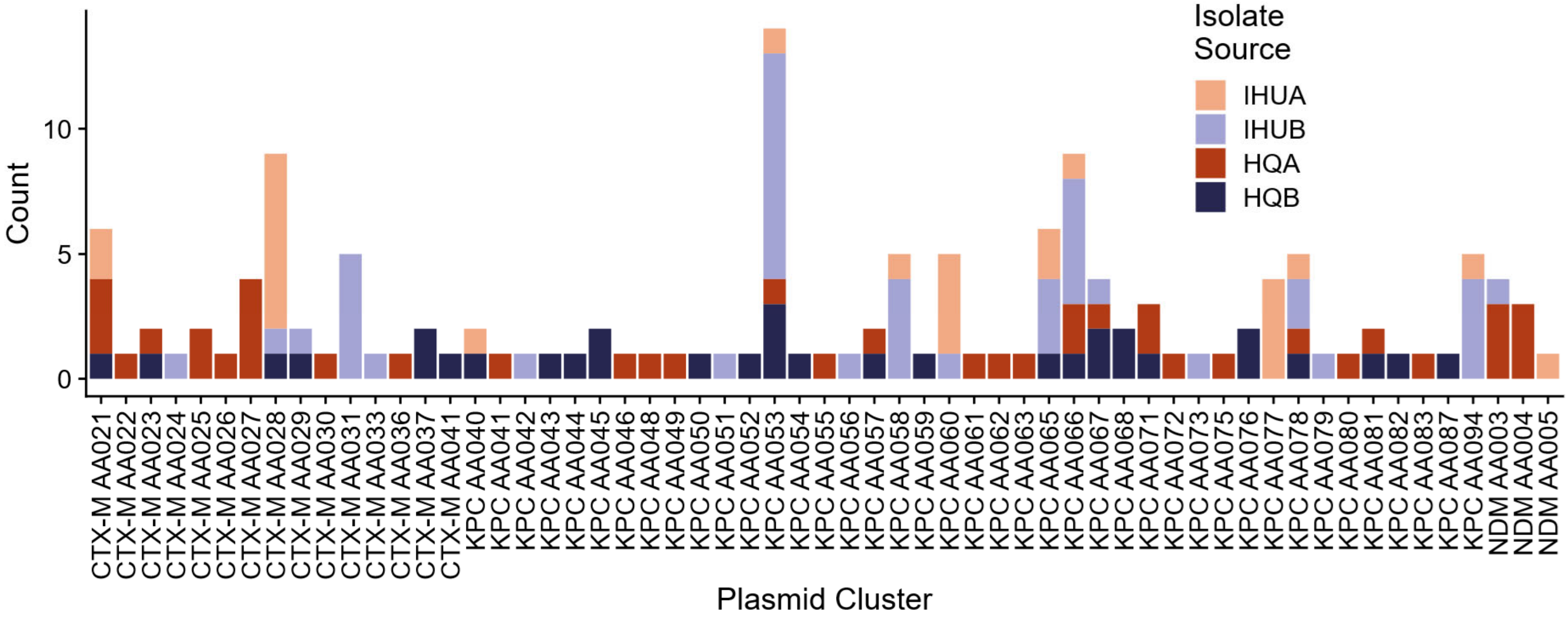


**Figure S8. Collection sources of ARG-containing plasmid clusters.** For each secondary cluster (Mash distance threshold 0.025) of blactxm, blaKPC, and blandm containing plasmids, bar plot heights indicate the number of total plasmids assigned to that cluster, with colors indicating isolate collection site (sink drains from inpatient hospital units IHUA or IHUB, or wastewater from hospital quadrants HQA or HQB). Plasmid sharing across sites was frequent; 3/15 (20%) $bla_{\text{CTX-M}}$-harboring secondary plasmid clusters, 6/41 (15%) $bla_{\text{KPC}}$ secondary clusters, and 1/3 (33%) $bla_{\text{NDM}}$ secondary clusters included plasmids found in both wastewater and sink drains.
