## Supplemental Figure 9 for "Interconnected reservoirs of multidrug-resistant bacteria and plasmids within hospital plumbing infrastructure"

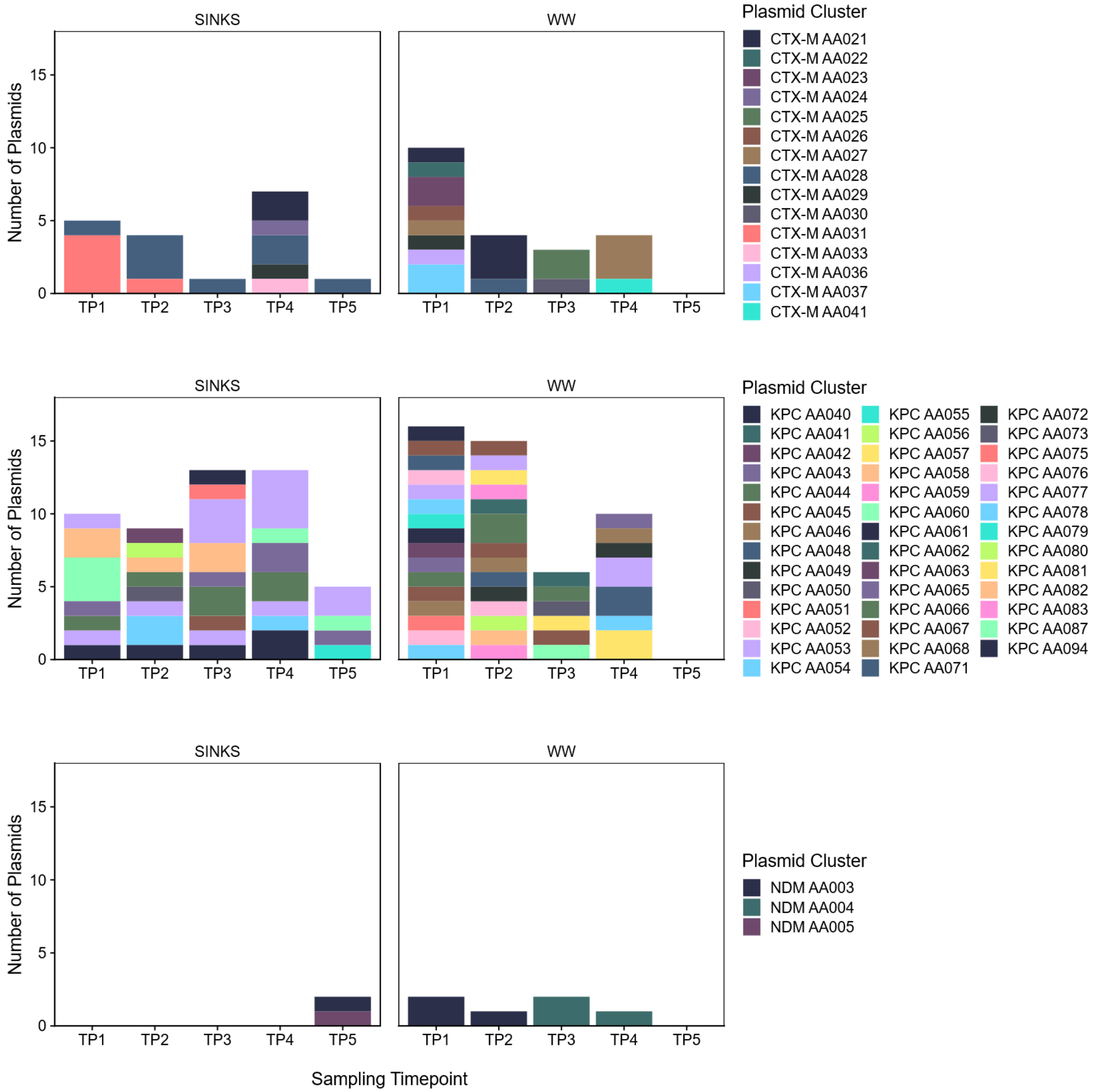


**Figure S9. Longitudinal prevalence of antibiotic resistance plasmid clusters.** Each row of panels displays the number of *bla*$_{CTX\text{-}M}$, *bla*$_{KPC}$, or *bla*$_{NDM}$ plasmids, respectively, belonging to a given secondary cluster (indicated by color; defined by a Mash distance threshold of 0.025) collected in sink drains (left column) or hospital wastewater (right column) at each sampling timepoint (TP1-TP5). Diverse *bla*$_{CTX\text{-}M}$ and *bla*$_{KPC}$ plasmids were identified in both environmental sources and across timepoints; however, sink drains tended to contain several larger plasmid clusters at a given timepoint while wastewater ARG-harboring plasmids were more evenly distributed across many secondary clusters. Comparison across sink drains and wastewater revealed that plasmid clusters were primarily constrained to a single environmental source, though at least one secondary cluster of plasmids harboring each of the three ARGs was shared between sources. Of the 15 total *bla*$_{CTX\text{-}M}$ plasmid secondary clusters, 4 (27%) were identified over at least two timepoints in the same environmental source, as were 14 out of 41 (34%) *bla*$_{KPC}$ and 2 out of 3 (67%) *bla*$_{NDM}$ secondary clusters. *bla*$_{CTX\text{-}M}$ plasmid cluster AA028 and 5 *bla*$_{KPC}$ clusters, KPC AA053, AA065, AA066, AA077, and AA094, were all highly persistent, identified in the same source at least four different timepoints. Of these, KPC AA053, AA066, and AA078 were notably persistent in both sink drains and wastewater.
