## Supplemental Figure 10 for "Interconnected reservoirs of multidrug-resistant bacteria and plasmids within hospital plumbing infrastructure"

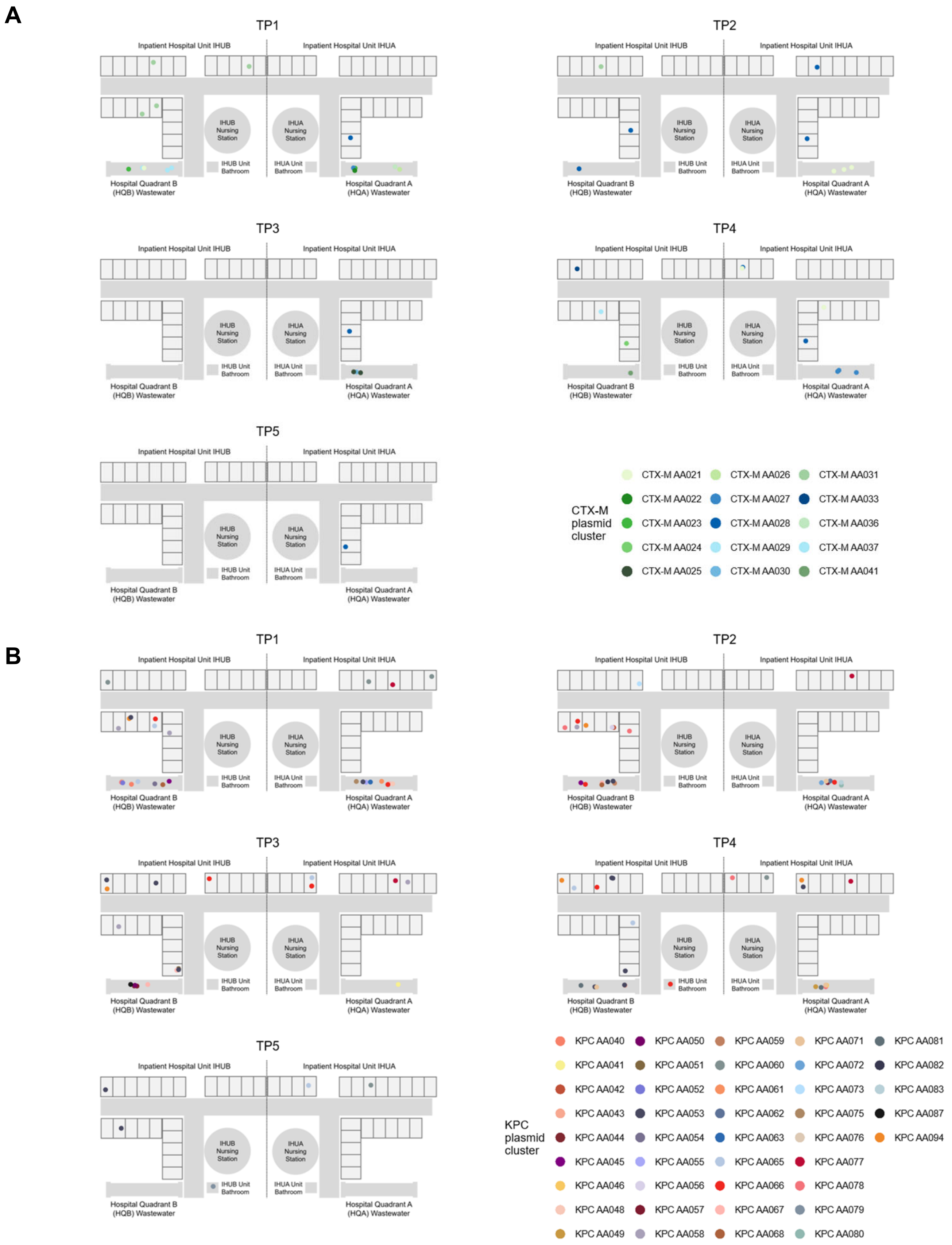


**Figure S10. Spatiotemporal analysis of persistent antibiotic resistance plasmids.** We identified 4 $\mathit{bla}_{\text{CTX-M}}$ and 14 $\mathit{bla}_{\text{KPC}}$ plasmid clusters (Mash distance < 0.025) which were present in the same environmental source (sink drains and/or wastewater) over at least two collection timepoints (Figure S8). For each of these persistent plasmid clusters, we plotted isolate collection sites (rooms where sink drains were located or hospital wastewater quadrant) to assess their spatial distribution across timepoints. We observed instances of persistence not only temporally, in the same source over multiple timepoints, but also spatially, in the same or nearby rooms (e.g., CTX-M AA028 in IHUA). We also identified several cases where the same plasmid cluster specifically and contemporaneously overlapped between inpatient hospital units and wastewater (e.g., KPC AA067 persistently identified in IHUB and HQB only, and not IHUA/HQA).
