## Supplemental Figure 11 for "Interconnected reservoirs of multidrug-resistant bacteria and plasmids within hospital plumbing infrastructure"

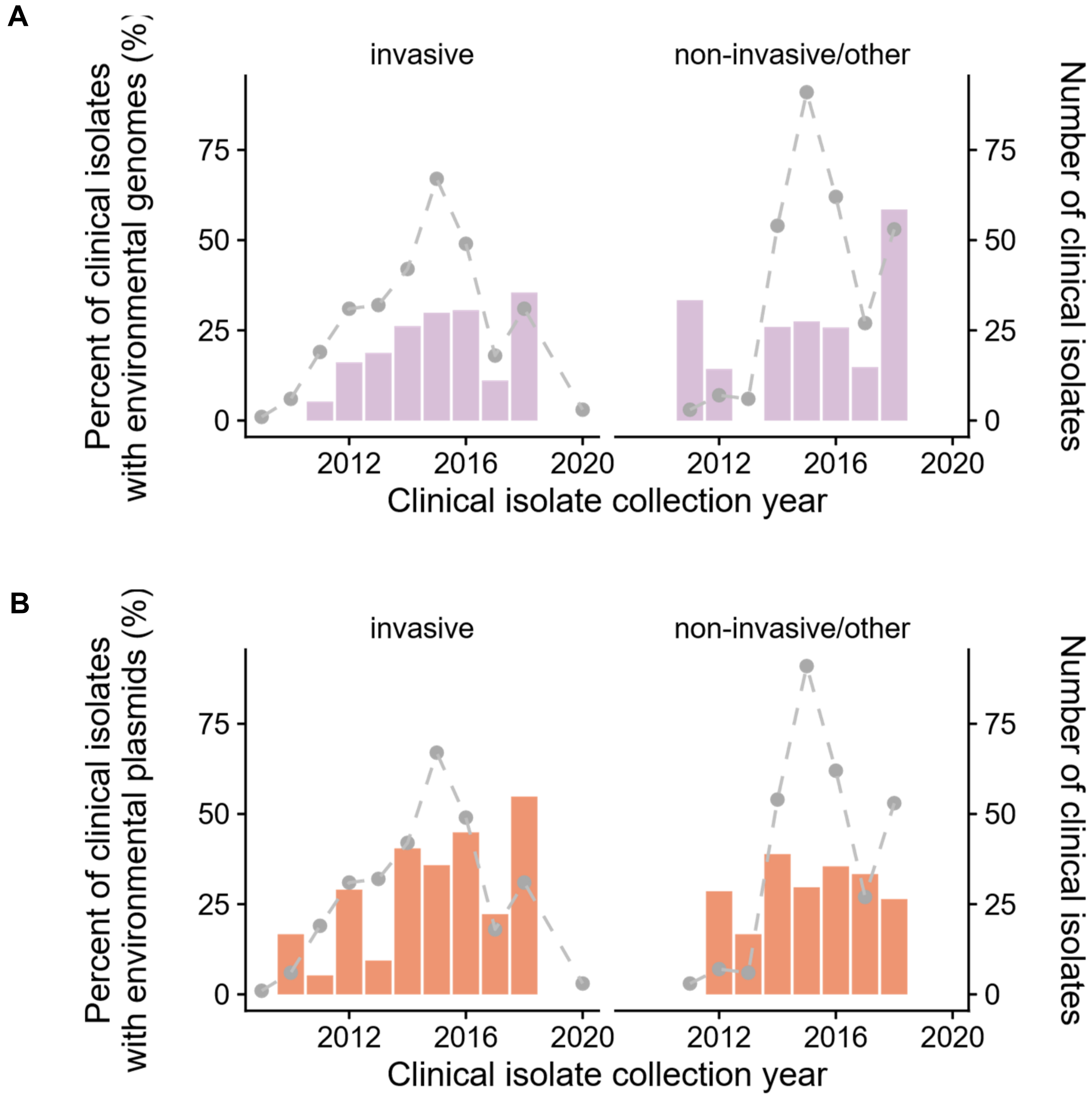


**Figure S11. Genome and antibiotic-resistant plasmid sharing between clinical and environmental isolates based on clinical collection source.** For all clinical isolates included in this study (n=602) from patient infections in 2009 through 2020, collection source was classified as invasive (blood, abdominal or intraabdominal, or respiratory cultures) or non-invasive (urine, wound, rectal swab, and other sites). Dashed lines and points in grey show the total number of invasive and non-invasive isolates included per year. Bar plots for each year indicate the proportion of clinical isolates which shared a genome (A) or plasmid (B) sequence with at least one environmental isolate collected from sink drains or hospital wastewater.
